# Monitoring the collective behavior of enzymatic nanomotors in vitro and in vivo by PET-CT

**DOI:** 10.1101/2020.06.22.146282

**Authors:** Ana C. Hortelao, Cristina Simó, Maria Guix, Sandra Guallar-Garrido, Esther Julián, Diana Vilela, Luka Rejc, Pedro Ramos-Cabrer, Unai Cossío, Vanessa Gómez-Vallejo, Tania Patiño, Jordi Llop, Samuel Sánchez

## Abstract

Enzyme powered nanomotors hold great potential for biomedical applications, as they show improved diffusion and navigation within biological environments using endogenous fuels. Yet, understanding their collective behavior and tracking them *in vivo* is paramount for their clinical translation. Here, we report on the *in vitro* and *in vivo* study of swarms of self-propelled enzyme-nanomotors and the effect of collective behavior on the nanomotors distribution within the bladder. For that purpose, mesoporous silica nanomotors were functionalized with urease enzymes and gold nanoparticles. Two radiolabeling strategies, i.e. absorption of ^124^I on gold nanoparticles and covalent attachment of an ^18^F-labeled prosthetic group to urease, were assayed. *In vitro* experiments using optical microscopy and positron emission tomography (PET) showed enhanced fluid mixing and collective migration of nanomotors in phantoms containing complex paths. Biodistribution studies after intravenous administration in mice confirmed the biocompatibility of the nanomotors at the administered dose, the suitability of PET to quantitatively track nanomotors *in vivo*, and the convenience of the ^18^F-labeling strategy. Furthermore, intravesical instillation of nanomotors within the bladder in the presence of urea resulted in a homogenous distribution after the entrance of fresh urine. Control experiments using BSA-coated nanoparticles or nanomotors in water resulted in sustained phase separation inside the bladder, demonstrating that the catalytic decomposition of urea can provide urease-nanomotors with active motion, convection and mixing capabilities in living reservoirs. This active collective dynamics, together with the medical imaging tracking, constitutes a key milestone and a step forward in the field of biomedical nanorobotics, paving the way towards their use in theranostic applications.

## Introduction

Self-propelled particles hold potential to overcome the biological barriers that limit current cancer nanomedicines, where only 0.7% of the administered dose reaches the target *in vivo*.(*1*) In this regard, micro- and nanomotors have demonstrated enhanced targeting properties(*2*–*5*) and superior drug delivery efficiency compared to passive particles.(*6*–*9*) Additionally, they outperform traditional nanoparticles in terms of penetration into biological material, such as mucus,(*10*–*13*) cells(*14*–*16*) or spheroids.(*4*, *17*) Particularly, using enzymes as biocatalysts is emerging as an elegant approach when designing self-propelled particles, due to the use of endogenous fuels, which enables nanomotors’ on-site activation and the design of fully biocompatible motor-fuel complexes. Moreover, the library of enzyme/substrate combinations permits the design of application-tailored enzymatic nanomotors, as is the case of urease-powered nanomotors for bladder cancer therapy.(*4*)

Yet, the large number of nanoparticles required to treat tumors(*18*) demands for a better understanding, control and visualization of nanoparticle swarms to aid in the evaluation of motile nanomedicines and facilitate the eventual translation into clinics. Indeed, collective phenomena commonly observed in nature (active filaments,(*19*) bacteria quorum sensing,(*20*) cell migration,(*21*) swarms of fish, ants, and birds)(*22*) also occurs in micro-/nanomotors, which demonstrated collective migration,(*23*–*26*) assembly,(*27*–*30*) or aggregation/diffusion behaviors *in vitro*.(*31*–*38*) *Ex vivo,* swarms of magnetic micropropellers demonstrated long-range propulsion through porcine eyes to the retina, suggesting potential as active ocular delivery devices.(*39*, *40*) *In vivo,* controlled swimming of micromotor swarms was shown in mouse peritoneal cavities, (*41*) and were tracked in the stomach(*24*) and intestines(*26*) in rodents using magnetic resonance imaging and photoacoustic computed tomography, respectively. These techniques, however, have low sensitivity and fail in providing quantitative information.

For this, positron emission tomography (PET), a non-invasive nuclear imaging technique widely used in clinics, is ideally suited. First, it is fully quantitative, and allows whole-body image acquisition. Second, it relies on gamma ray detection, which have no tissue penetration limit, turning this imaging modality into a fully translational tool. Finally, it is extremely sensitive, thus providing good quality images by administering sub-pharmacological dosages of the radiolabeled entity.(*42*, *43*) Surprisingly, its application for *in vivo* tracking of micro-/nanomotors has barely been explored.(*44*)

Herein, we investigate the swarm behavior of enzyme-powered nanomotors *in vitro* and *in vivo,* using PET in combination with computed tomography (CT). We prepared enzyme-powered nanomotors based on a FDA-approved mesoporous silica chassis(*45*) and labeled them with Iodine-124 (^124^I; half-life = 4.2 days) and Fluorine-18 (^18^F; half-life = 119.7 minutes). The suitability of PET imaging to investigate the swarm behavior of labeled nanomotors in the presence of the fuel was first demonstrated *in vitro* using tailored phantoms. Stability of the label and whole-body biodistribution was then investigated *in vivo* after intravenous administration into mice. Finally, time-resolved evidence of the motile properties of the nanomotors in the bladder was obtained after intravesical instillation.

## Results

Nanomotors were prepared by synthesizing mesoporous silica nanoparticles (MSNPs) using a modification of the Stöber method (see experimental section for details),(*46*) and their surface was modified with amine groups by attaching aminopropyltriethoxysilane (APTES).(*47*) The amine groups were subsequently activated with glutaraldehyde (GA) to enable the covalent binding of the enzymes and the heterobifunctional polyethylene glycol (NH_2_-PEG-SH). Finally, AuNPs were anchored to the surface of the nanomotors by attachment to NH_2_-PEG-SH *via* thiol-gold chemistry (Figure 1a).

**Figure 1.**
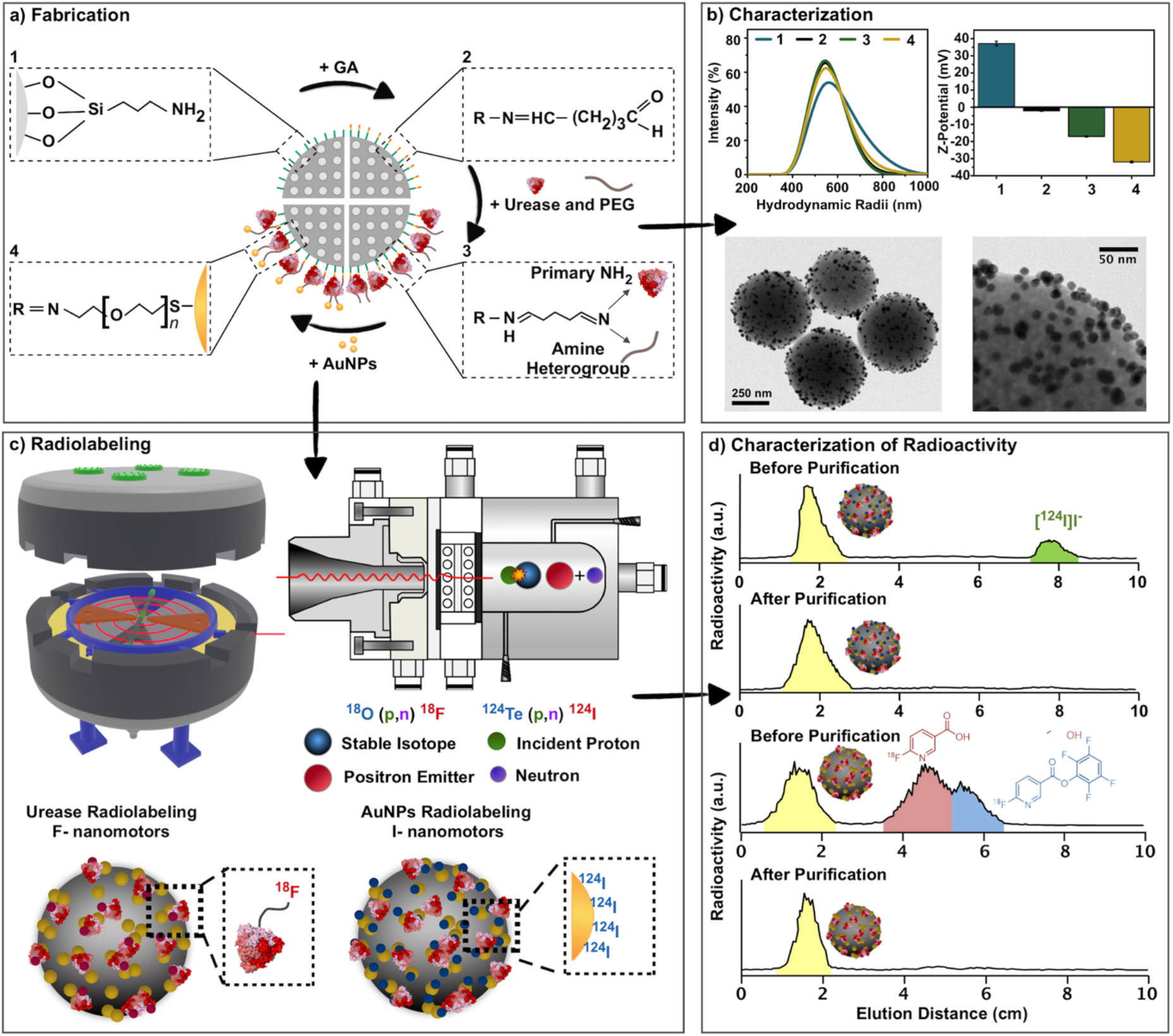
Preparation and characterization of radiolabeled urease-powered, AuNP-decorated nanomotors. a) Process flow of nanomotors fabrication steps. b) Characterization of the nanomotors by dynamic light scattering (DLS), Z-potential and transmission electron microscopy (TEM). c) Nanomotors radiolabeling using either Fluorine-18 or Iodine-124, to yield ^18^F-nanomotors or ^124^I-nanomotors, respectively; d) Labeling efficiency and radiochemical purity monitored by instant thin layer chromatography (iTLC).

The hydrodynamic radii and stability of nanoparticles along the different fabrication steps were monitored by dynamic light scattering (DLS) analysis. The single population peak corresponding to MSNPs became sharper upon addition of NH_2_-PEG-SH, suggesting that PEG provides steric stabilization to the nanomotors in solution.(*48*) The single population peak was conserved after attachment of AuNPs, evidencing that the final synthetic step of the process did not induce aggregation.

Electrophoretic mobility analysis was performed to characterize the surface properties of the particles after each functionalization step. Z-Potential values for MSNP-NH_2_ were found to be 37.1 ± 1.4 mV. The trend towards negatively charged surface in subsequent synthetic steps confirms the successful activation of the MSNP-NH_2_ with GA (Z-potential = −2.2 ± 0.2 mV), and incorporation of NH_2_-PEG-SH and urease enzymes (Z-potential = −32.0 ± 0.6 mV). The final negative value can be explained by urease’s isoelectric point (*ca.* 5), which indicates that at pH > 5 the net charge of urease is negative, and the negative charge of the AuNPs.(*49*) The spherical nanoparticles had an average diameter of 507.8 ± 3.4 nm and a stochastic distribution of the AuNPs on the surface, as shown in the TEM image (Figure 1b and Figure S1).

Next, to enable the visualization of the nanomotors by PET, we radiolabeled them with the positron emitters ^18^F and ^124^I (Figure 1c). For the radiofluorination, a straightforward strategy based on the use of the pre-labeled prosthetic group [^18^F]FPyTFP was developed.(*50*) Taking advantage of the free amino groups on the enzyme, sufficient labeling yield (30% with respect to [^18^F]FPyTFP, decay-corrected) was achieved by incubation (35 minutes at room temperature) of the nanomotors with the prosthetic group. Radioiodination was achieved by direct absorption of ^124^I on AuNPs on the nanomotors.(*51*) The radiochemical yield was 71 ± 2 %, due to the high affinity binding between gold and iodine. Radiochemical purity after purification was ≥99 % for both cases, as determined by instant thin layer chromatography (Figure 1d).

The self-propulsion of urease-modified micro-/nanomotors(*4*, *52*–*54*) is caused the asymmetric release of ionic species from the particles to the solution(*55*) which stems from the catalytic decomposition of urea into carbon dioxide and ammonia. Despite the recent advances in the field, enzyme nanomotors have been only studied from single particle to a few particles case, being their collective swarming behaviour not investigated to date.

The motion dynamics and swarming behavior of nanomotors *in vitro* was first investigated by optical microscopy. A 2 μL droplet of nanomotors suspension was placed onto a Petri dish containing either phosphate buffer saline (PBS) or a 300 mM urea solution, since urea concentrations in the bladder can reach up to this concentration,(*56*) and videos were recorded over 2 minutes (Figures 2a, 2b and S2 and video S1). Figure 2a suggests that after addition of the droplet in PBS, the nanomotors stay at the seeding point following a stochastic distribution, which remains unaltered over time as observed in the histograms of pixel intensity distributions obtained for the selected ROI (Figure 2c). Preferential tracks are observed immediately after seeding due to the motion of the nanomotors in urea, resulting in an anisotropic distribution of the nanomotors throughout the dish which evolves over time, as observed in Figure 2b and evidenced by the dynamic changes in the histograms (Figure 2d).

**Figure 2.**
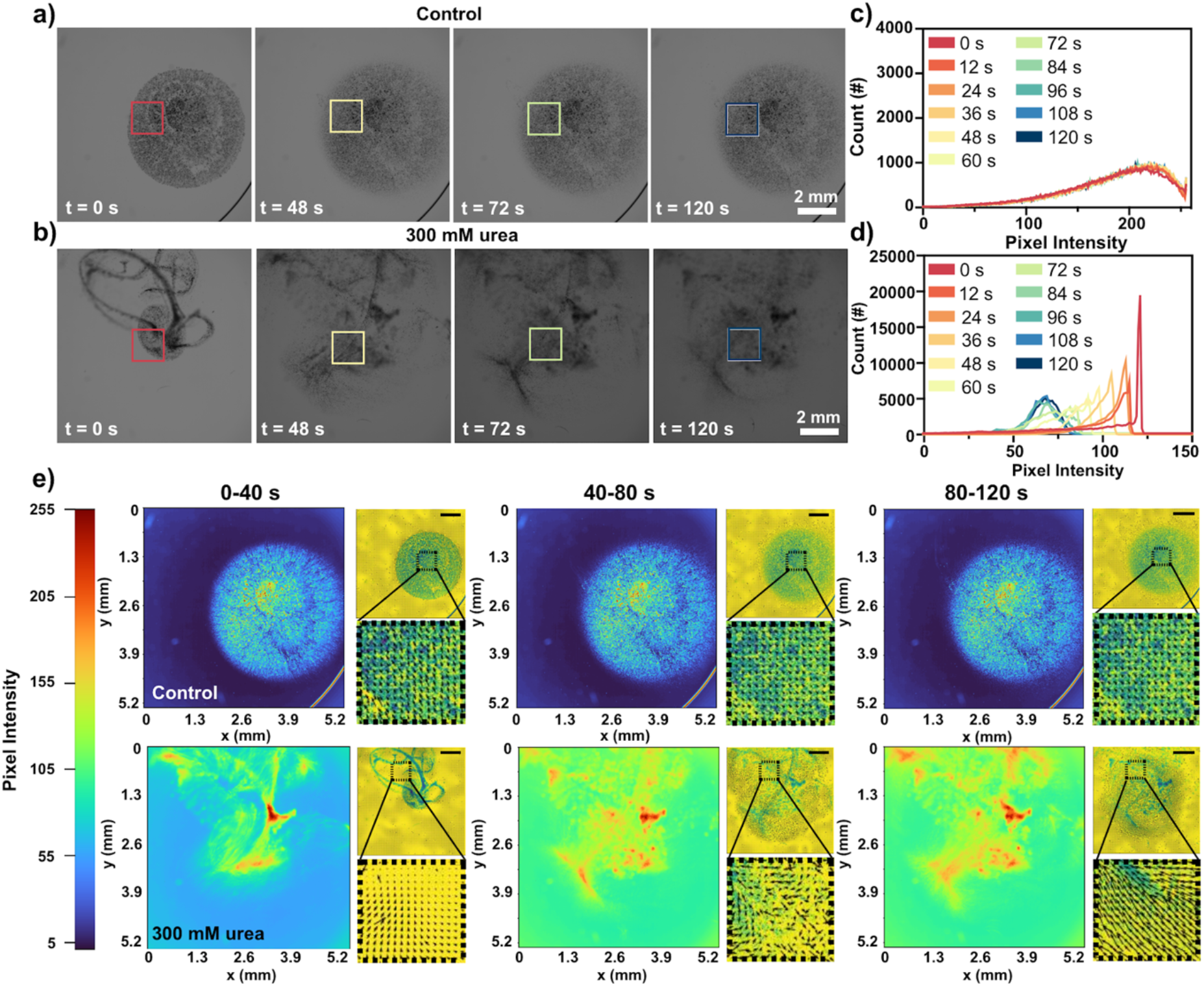
*In vitro* imaging of urease-powered nanomotor swarms by optical microscopy. Snapshots of a population of urease-powered nanomotors in a) PBS and b) 300 mM urea in PBS. Corresponding pixel intensity distribution histograms within the regions of interest noted in the images of nanomotors in c) PBS and d) 300 mM urea in PBS. Density maps (large panels) obtained by calculating the sum pixel intensity over periods of 40 seconds of the videos and particle image velocimetry (PIV) (small panels and zoom out) of e) nanomotors in PBS (top) and in 300 mM urea in PBS (bottom); scale bar in small panels is 1 mm.

To gain further insight into swarm dynamics, we then generated density maps. These were obtained by representing sum pixel intensity values of video frames within pre-established time ranges (0-40 s, 40-80 s and 80-120 s) using a colormap (Figure 2e and S3). In absence of urea, density maps did not significantly vary with time, and pixel intensity values for most of the population were around 60 (Figure 2e, top). On the other hand, the density maps for nanomotors in urea show multiple paths followed by the nanomotors immediately after droplet addition (Figure 2e, bottom). This is shown as a fast change in the density maps color, where the intensity of the overall field of view and nanomotor flocks rapidly increases, which indicates high mobility and low residence time.

To investigate if the activity of the nanomotors and the swarming behavior were associated with fluid flow generation, we carried out particle image velocimetry (PIV) analyses on the optical microscopy videos (Figure 2e, inserts), PIV is an optical technique that allows the visualization of fluid flow associated with particle motility, indicating fluid and particle displacements in the form of vector fields. In the control condition (Figure 2e, top), the quiver plots show velocity vectors with low magnitude and in scarce amount (small panels). Oppositely, the quiver plots obtained for nanomotors in urea show dispersed vector fields emerging not only from the swarms but also from the surrounding fluid. The evolution of the vector fields in time denotes the formation of vortices and fronts that promptly dissociate. Altogether, these results demonstrate that the activity of the nanomotors and their emergent collective behavior induce enhanced convection and fluid mixing, similar to what has been observed with catalytic micromotors.(*57*–*59*)

To confirm these results and assess the feasibility of PET imaging to track swarm dynamics, we carried out parallel experiments with radiolabeled nanomotors. PET images obtained at different times after droplet addition confirmed that the nanomotors remain at the seeding point in water (Figure 3a and video S2), and uniformly distribute over the whole volume of the Petri dish in the presence of urea (Figure 3b). This behavior could be also clearly visualized in 3D histograms representing the concentration of radioactivity throughout the petri dish (Figures 3c and 3d). Overall, these experiments demonstrated the suitability of PET imaging to obtain time-resolved quantitative information about the nanomotors swarming dynamics *in vitro*.

**Figure 3.**
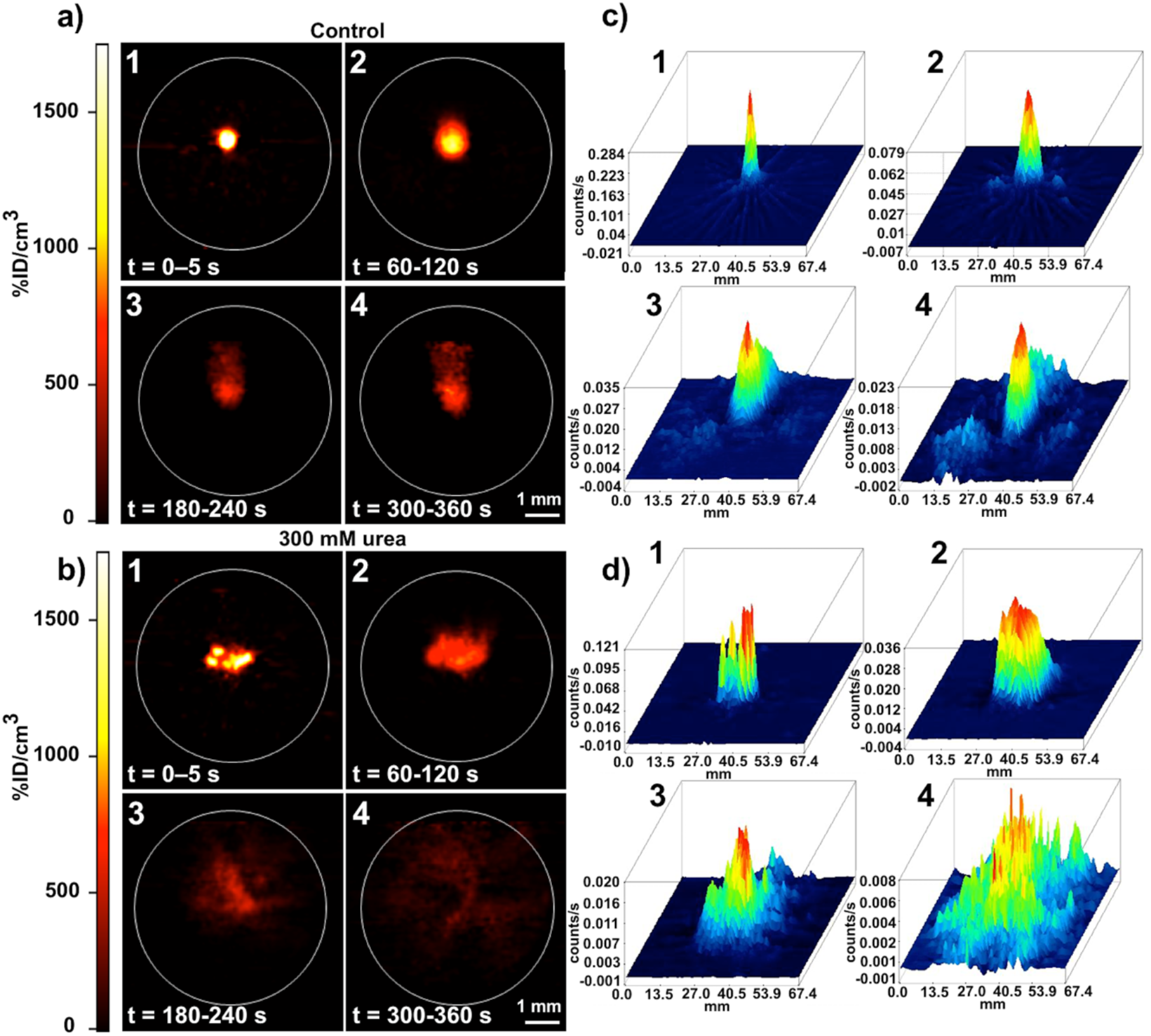
*In vitro* imaging of urease-powered ^18^F-nanomotor swarms by PET. Snapshots of a population of urease-powered nanomotors in a) water and b) 300 mM urea imaged by PET. Pixel intensity distribution histograms of the population of nanomotors in the field of view in c) water and d) 300 mM urea.

The *in vitro* swarm behavior of the nanomotors was investigated in four polydimethylsiloxane phantoms with increasing degrees of complexity (Figure 4a top; 4b-e; and S4). The selected phantoms comprise different path shapes: (*i*) straight, (*ii*) rectangular, (*iii*) curved, and (*iv*) a curved path with longer straight trenches. In each phantom, one channel was filled with water and another channel with 300 mM urea solution in water. The nanomotors (either ^124^I- or ^18^F-labeled) were seeded at the edge of each channel and dynamic PET images were immediately acquired for 25 minutes.

**Figure 4.**
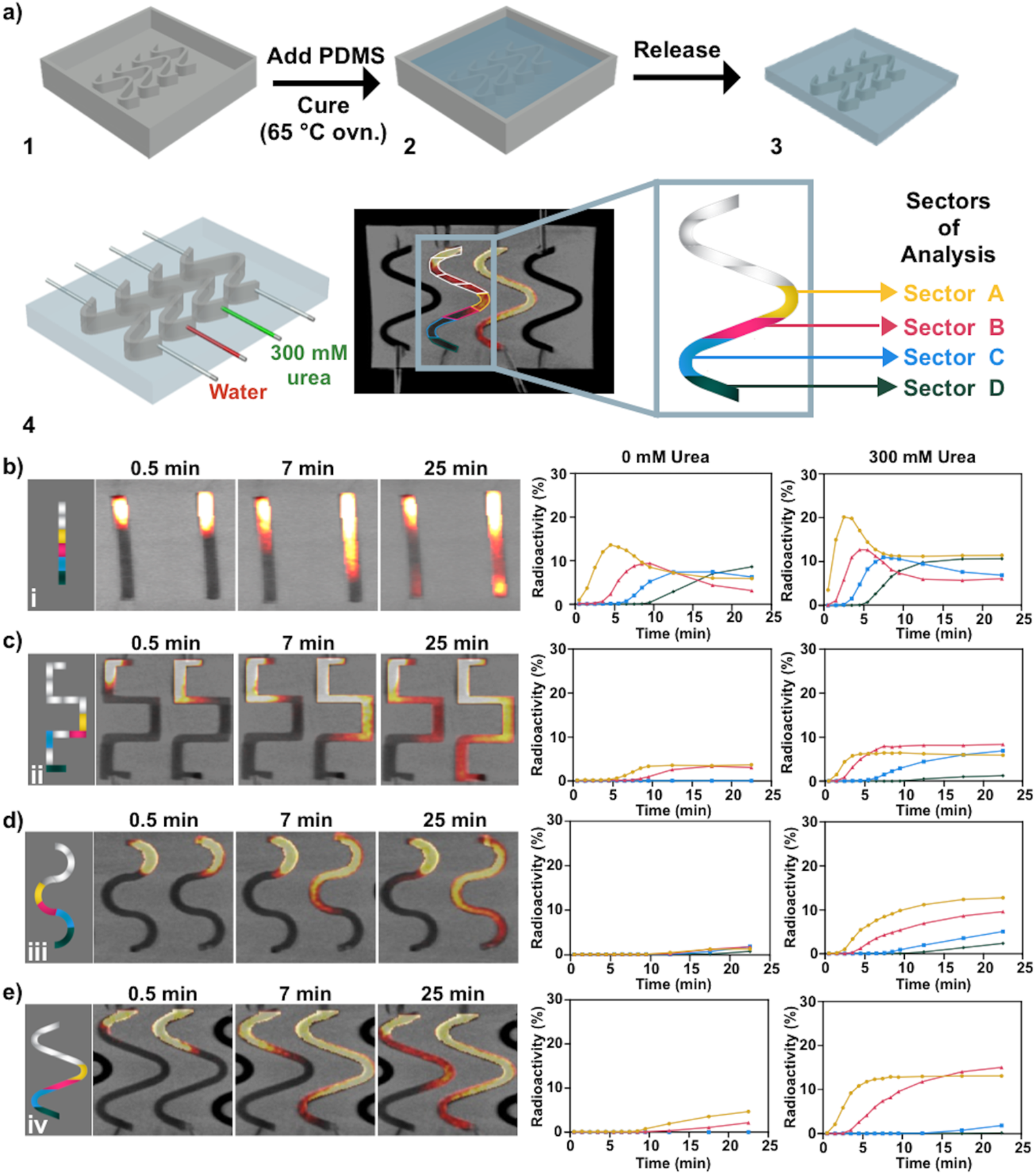
Effect of complex paths on the *in vitro* motion of ^18^F radiolabeled urease-powered nanomotors studied by PET-CT imaging; a) scheme depicting the fabrication process of 3D phantoms comprising different complex geometries (steps 1 to 4) and the corresponding method of analysis; b–e) on the left, PET images (coronal projections) obtained at 0.5, 7 and 25 minutes after seeding of the nanomotors in different-shaped phantoms. Scale bars are 3 mm. For each phantom, one of the channels was filled with water (left) and another channel was filled with 300 mM urea solution (right). Quantitative results of the normalized concentration of radioactivity for each sector as a function of time are shown on the right.

The images obtained at *t* = 0.5, 7 and 25 minutes after seeding ^18^F-nanomotors (Figure 4b-4e and video S3) prove that urea has a prominent effect on the motion dynamics of the nanomotors throughout the paths. Irrespective of the path shape, the nanomotors reached the end of the channel at *t* = 25 minutes in the presence of the fuel, while most nanomotors remained close to the seeding point in water and only minimal movement could be detected.

In order to get quantitative data, dynamic PET images were analyzed by dividing the phantoms in sectors (Figure 4a, bottom-right) and the concentration of radioactivity in each sector (normalized to the total amount of radioactivity in the channel) was determined as a function of time. Differences were evident even in the less restrictive phantom (straight shape), in which free diffusion of non-activated nanomotors is more favored. Curves with higher slope were obtained in sectors A-D in urea (Figure 4b), confirming that the speed at which the nanomotors reach the different sectors is higher than in water. In addition, the fraction of nanomotors that reached the second half of the phantom (sectors B-D) at *t*=25 minutes was higher in the presence of urea (24%) than in water (18%). A similar trend was observed for ^124^I-nanomotors (Figure S5).

These differences increased when the mobility was limited by introducing complex paths (phantoms ii-iv). Figure 4c-4e show that nanomotors reach sectors B-D much faster in urea than in water, which was confirmed by the amount of nanomotors reaching the second half of the phantom at the end of the imaging session. Values of 34.2, 17.1 and 17.0% were obtained for phantoms ii-iv in urea, while values in water were as low as 8.5, 4.2 and 2.1%. Equivalent results were obtained for ^124^I-nanomotors (Figure S5), where the fraction of radioactivity in the second half of the phantom at the end of the study was 7.4, 4.2 and 2.0% for phantoms ii-iv in water, while values in urea were 38.2, 20.6 and 27.0%, respectively. Small differences in the values obtained using the two radionuclides might be due to inherent limitations of PET imaging. Indeed, the diameter of the channels is very close to the spatial resolution of our PET imaging system, which is 1.2 mm full width half maximum for ^18^F, and significantly lower for ^124^I due to less favorable emission properties.(*60*) This, together with the presence of a high concentration of radioactivity in small volumes, can lead to partial volume effect, thus causing some error in the absolute quantification values. In spite of this limitation, PET imaging results unambiguously confirmed the enhanced mobility of enzyme-powered nanomotors in urea, with an effect that becomes more relevant when the complexity of the path increases.

A biodistribution study of ^18^F-nanomotors and ^124^I-nanomotors in female mice after intravenous administration was carried out in order to (i) demonstrate the suitability of *in vivo* PET imaging to quantitatively track the nanomotors at the whole body level; and (ii) evaluate their radiochemical stability *in vivo*.

Images obtained after administration of ^18^F-nanomotors (Figure 5a and video S4) evidence a biodistribution profile with initial accumulation in the lungs and the liver, and progressive elimination of the radioactivity *via* urine, confirmed by image quantification (Figures 5b and 5c). These results suggest rapid uptake by the mononuclear phagocyte system, as typically observed with intravenously administered nanoparticles of this size.(*61*) Since the nanomotors are above the estimated size threshold for glomerular filtration (*ca.* 8 nm),(*62*) the increase in radioactivity in urine indicates a slow detachment of the radiolabel from the nanomotors.

**Figure 5.**
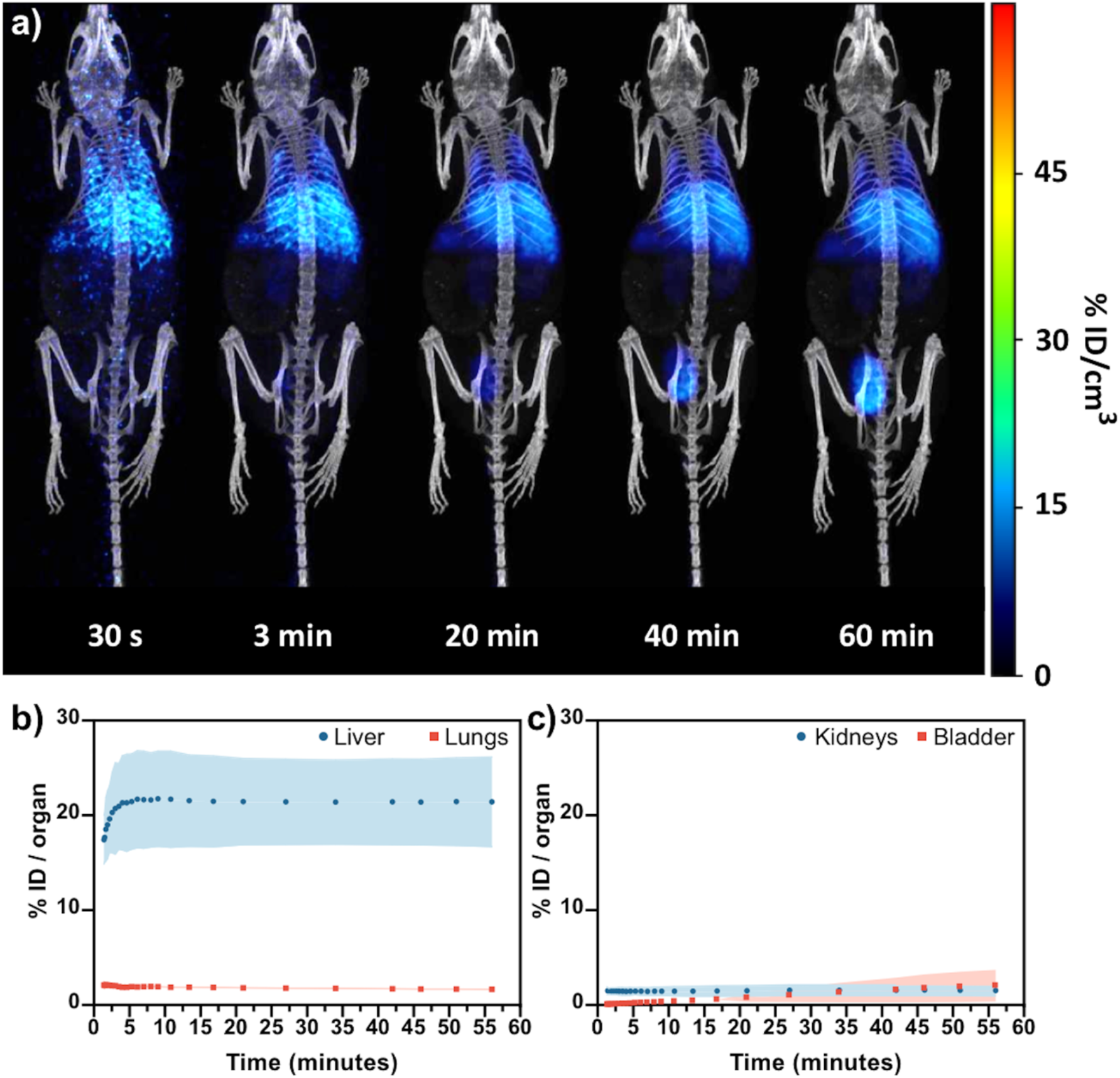
Analysis of the biodistribution of ^18^F-labelled urease-AuNP nanomotors injected intravenously in female mice. a) PET-CT images (maximum intensity projections, coronal views) obtained at different time points after intravenous administration of ^18^F-labeled urease AuNP-nanomotors; b) time-activity curves in the liver and lungs and (c) kidneys and bladder, as determined by PET imaging. Results are expressed as % of injected dose per organ (mean ± standard deviation, N = 2).

^124^I-nanomotors also show initial accumulation in the liver and the lungs (Figure S6). Strikingly, the concentration of radioactivity in both organs progressively decreases afterwards, paralleled by an increase in the thyroid gland, stomach and urine (Figure S6 b-d). The thyroid gland and urine are the metabolic sites of free iodine, and hence this suggests fast nanomotor deiodination, explained by the desorption of ^124^I from the gold surface.

Altogether, these results demonstrate that positron emitter-labeled nanomotors can be tracked *in vivo* after administration into living organisms using PET, and that ^18^F radiolabeling yields higher radiochemical stability *in vivo* than ^124^I-radiolabeling, since the detachment of the radiolabel is almost negligible over the duration of the study. Importantly, no adverse effects were observed in the animals for 2 weeks following imaging sessions, thus suggesting that the administered dose is below the maximum tolerated dose.

Envisioning bladder cancer imaging and therapy, and since the nanomotors size prevents accumulation in the bladder after intravenous injection, we studied their behavior after intravesical instillation. This administration route is well established in bladder cancer therapy, since it maximizes the concentration of the drug in the target organ, resulting in improved efficacy and fewer side-effects.(*63*)

Owing to the stability of the ^18^F radiolabel demonstrated in the biodistribution study, only ^18^F-nanomotors were used in these experiments. In addition, the use of ^18^F should facilitate clinical translation, as it can be easily produced in biomedical cyclotrons and presents superior physical properties. When ^18^F-nanomotors were intravesically instilled using 300 mM urea in water as the vehicle, we observed a uniform distribution of the radioactivity immediately after instillation, followed by a two-phase formation at *t* = 15 minutes (Figure 6b, 1 and video S5). Unexpectedly, a progressive cancelation of the difference between the two phases was observed at longer times, leading to a uniform distribution of radioactivity at *t* = 45 minutes. On the contrary, when administered in water, the phase separation was maintained (Figure 6b, 2). These observations were further confirmed by analysis of the concentration of radioactivity in two volumes of interest (VOIs) drawn within the two phases observed. Indeed, for ^18^F-nanomotors injected in urea, the time activity profile (Figure 6c, 1) shows that the concentration of radioactivity in both regions is close to 50% immediately after instillation (homogeneous distribution). The value in VOI 1 progressively increases to reach a maximum at ca. 1000 s, and decreases afterwards to recover starting values at *t* > 2000 s, confirming that the concentration of radioactivity within the bladder had regained homogeneity. In contrast, time activity curves obtained under the control condition show a progressively divergent trend (Figure 6c, 2), confirming that the concentration of radioactivity in the bladder is not homogeneous by the end of the study. We hypothesize that immediately after nanomotors instillation in the empty bladders, fresh urine starts entering. This (radioactivity-free) urine displaces the solution present in the bladder (containing the labeled nanosystems) and two different phases are formed. Since ^18^F-nanomotor swarms can actively move and enhance mixing in urea, they reverse the phase separation. However, when instilled in water, the separation of the two phases prevents the nanomotors being in close contact with the urea (present only in the fresh urine) and consequently nanomotors lack motility, thus maintaining the phase separation. The observed phenomenon is potentially advantageous in the design of active drug delivery systems, where homogeneous distribution of the delivery vehicles is required to ensure reaching the target site.

**Figure 6.**
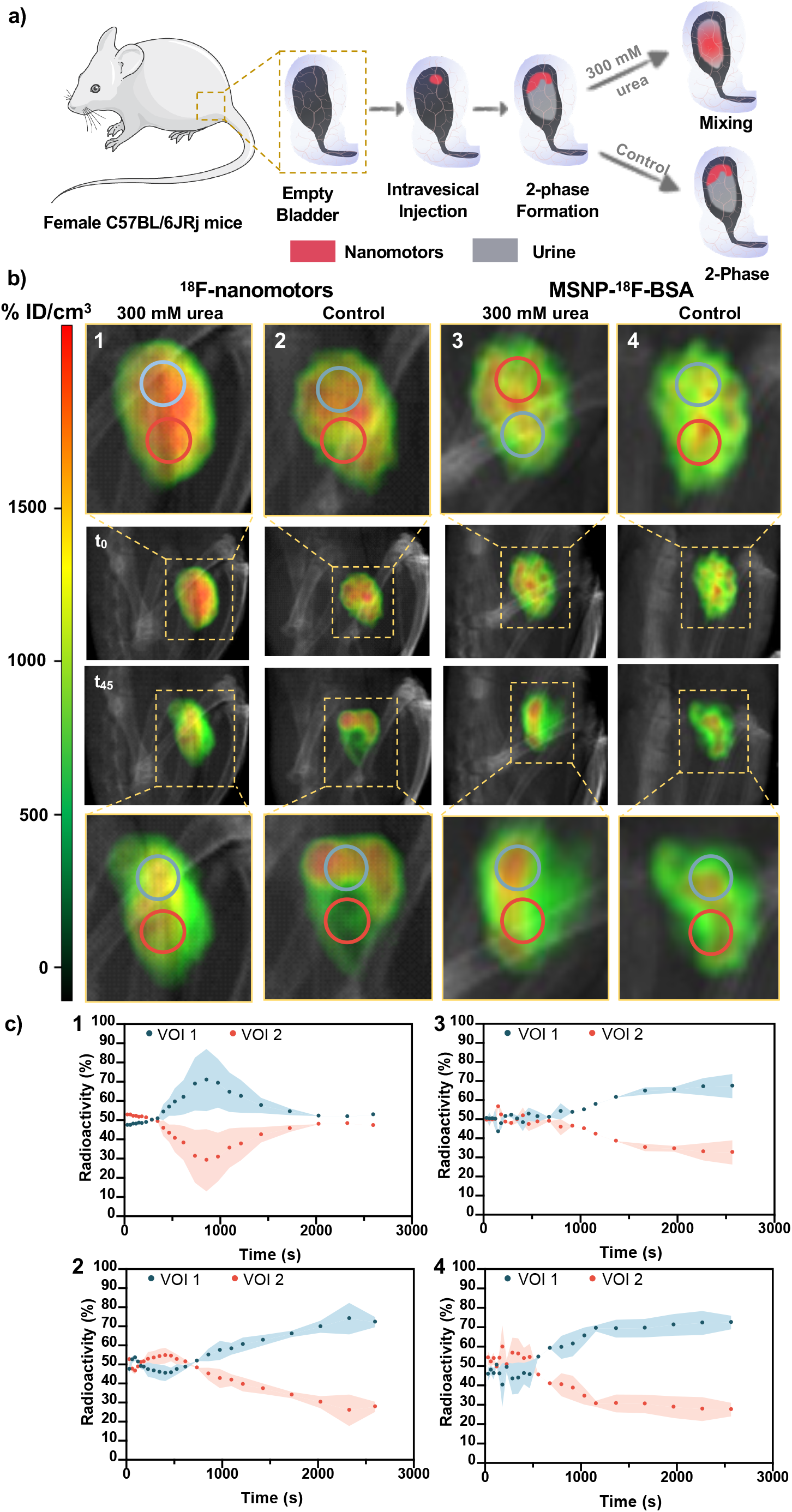
PET-CT analysis of the biodistribution in the bladder of ^18^F-nanomotors and MSNP-^18^F-BSA administered via intravesical instillation. a) Scheme depicting the administration method and phenomena observed. b) PET-CT images obtained at different time points after the intravesical instillation of ^18^F-labeled urease nanomotors in urea (1) and water (2), and control BSA-^18^F-MSNP in urea (3) and water (4). c) Quantification analysis of the PET-CT images obtained.

As an additional proof of our hypothesis, we administered ^18^F-labeled BSA modified particles (MSNP-^18^F-BSA), which do not show self-propelling properties neither in urea nor water. In this case, time activity curves followed the same trend as for ^18^F-nanomotors instilled in water (Figure 6b and 6c, *3* and *4*), confirming that the lack of motility of the particles results in non-homogenization of the two phases formed in the bladder cavity.

## Conclusions

We designed AuNPs-decorated enzyme powered nanomotors and investigated their swarming behavior *in vitro,* both using optical microscopy and by PET-CT. When swimming without boundaries, we observed that the nanomotor swarms lead to the formation of vortices and unstable fronts, and that this emergent collective behavior induces enhanced fluid convection and mixing. Moreover, when the nanomotors are subjected to boundaries in the form of complex geometries with increasing degrees of complexity, the collective motion allows them to overcome the hurdles encountered, *i.e.* the turns and angles present on the paths and reach the end of the track.

We studied the suitability of PET-CT to image nanomotors *in vivo* and at the whole-body level, using two different radiolabeling strategies: 1) attach ^124^I isotope to the AuNPs and 2) directly label the enzymes with ^18^F isotope. We demonstrated not only that PET-CT is a highly adequate technique to image nanomotors, but also that the direct label of enzymes affords higher radiolabeling yields and a more radiochemically stable structure.

Envisioning the application of urease-powered nanomotors in the treatment of bladder diseases, we studied their motility within mice’s bladder, verifying that the swarming behavior and dynamic distribution of enzymatic nanomotors also occurred *in vivo*. When intravesically instilled in mice, the nanomotors were able to move in swarms and exhibit enhanced fluid mixing, which led to their homogenous distribution in the whole bladder cavity. In contrast, passive particles resulted in a non-homogeneous distribution, where a 2-phase distribution was observed.

In conclusion, we studied the collective behavior of enzymatic nanomotors *in vitro* and *in vivo* by a combination of optical microscopy and PET-CT imaging techniques. This work proves that PET-CT imaging is a valuable technique for the tracking nanomotor swarms both *in vitro* and *in vivo*. The swarming behavior and fluid mixing of enzyme nanobots observed upon intravesical injection holds a great potential towards biomedical applications, where their active homogeneous distribution and enhanced fluid mixing could be exploited for targeting and drug delivery purposes.

## Materials and Methods

### Materials

Ethanol (99%), methanol (99%), hydrochloric acid (37% in water), ammonium hydroxide (25% in water), tetraethylorthosilicate (TEOS, 99%), triethanolamine (TEOA, 99%), cetyltrimethylammonium bromide (CTAB, 99%), 3-aminopropyltriethoxysilane (APTES, 99%), glutaraldehyde (GA, 25% in water), urease (from Canavalia ensiformis, type IX, powder, 50 000− 100 000 units per gram of solid), urea (99.9%), bovine serum albumin (lyophilized powder), and HS-PEG5K-NH2 (HCl salt) were purchased from Sigma-Aldrich. Phosphate-buffered saline (PBS) were purchased from Thermo Fisher Scientific. Standard Grey V4 resin (FLGPGR04) for the 3D printing of rigid molds was purchased at FormLabs. SYLGARD™ 184 Silicone Elastomer was purchased from Ellsworth Adhesives Ibérica.

### Instruments

Transmission electron microscopy (TEM) images were captured using a JEOL JEM-2100 microscope. Hydrodynamic radii and electrophoretic mobility measurements were performed using a Wyatt Möbius. Optical videos were recorded using an inverted optical microscope (Leica DMi8) equipped with a 2.5x objective. The molds for the phantoms were fabricated by using a Stereolitography (SLA) 3D printer FormLabs 2 (FormLabs Inc.). The fluorine radionucleotide was synthesized using a TRACERlab FX-FN from GE Healthcare. The molecular imaging experiments were performed using a MOLECUBES β-CUBE (PET) and the MOLECUBES X-CUBE (CT) scanner.

### Animals

Female mice (C57BL/6JRj, 8 weeks, Janvier lab; 16 animals, see below for number of animals under different experimental scenarios) weighing 20 ± 3 g were used to conduct the biodistribution studies. The animals were maintained and handled in accordance with the Guidelines for Accommodation and Care of Animals (European Convention for the Protection of Vertebrate Animals Used for Experimental and Other Scientific Purposes) and internal guidelines. All experimental procedures were approved by the ethical committee and the local authorities before conducting experimental work (Code: PRO-AE-SS-059).

### Synthesis of Mesoporous Silica Nanoparticles (MSNP)

The MSNPs that serve as chassis for the fabrication of the nanomotors were synthesized by sol-gel chemistry using a modification of the Stober method.(*46*) Briefly, a mixture of TEOA (35 g), milli-Q water (20 mL) and CTAB (570 mg) was placed in a 3-mouthed round bottom flask and heated up to 95°C in a silicon oil bath, under reflux and stirring, for 30 minutes. After this, TEOS (1.5 mL) was added dropwise. The reaction took place for 2 hours, under stirring and reflux at 95°C and then the resulting MSNPs were collected by centrifugation (3 times, 1350g, 5 min). The CTAB was then removed by reflux in acidic methanol. For this, the MSNPs were suspended in a methanol (30 mL) and hydrochloric acid (1.8 mL) mixture, placed in a one-mouthed round bottom flask in a silicon oil bath at 80°C and refluxed for 24 hours. Finally, the MSNPs were collected by centrifugation and washed thrice in ethanol and thrice in milli-Q water (3 times, 1350g, 5 min). The concentration of the final dispersion was evaluated by dry-weighing.

### Amine modification of MSNPs

The surface of the MSNPs was then modified to carry free amine groups, using a modification of a reported method.(*47*) Briefly, a suspension of MSNPs (2 mg/mL) in water was placed in a round bottom flask, and heated up to 50 °C under vigorous stirring. Then, APTES was added to the dispersion to a final concentration of 5 mM. The reaction took place under reflux, at 50°C, for 24 hours, after which the MSNPs-NH_2_ were collected and washed thrice in water by centrifugation (3 times, 1350g, 5 min). The concentration of the final dispersion was evaluated by dry-weighing.

### Synthesis of Gold Nanoparticles (AuNPs)

The AuNPs were synthesized according to a previously reported method.(*64*, *65*) Briefly, all the necessary material was cleaned using freshly prepared *aqua regia,* then rinsed extensively with water and dried in air. Then a solution of 1 mM of AuCl_4_ was heated up to boil under stirring, in a round bottom flask integrated in a reflux system. After this, 10 mL of a NaCit solution (30.8 mM) was added and the solution was boiled for 20 minutes, turning red in color. The solution was stirred without heating for 1 hour, reaching room temperature. The resulting AuNPs dispersion was stored at room temperature in the dark. The Z-potential of the synthesized AuNPs in water was −40.26 ± 2.23 mV and their hydrodynamic radii 10.4 ± 0.1 nm (n = 10).

### Fabrication of Nanomotors

The MSNP-NH_2_ were re-suspended in PBS 1- at a concentration of 1 mg/mL and a total volume of 900 μL and activated with GA (100 μL). The reaction took place for 2.5 hours, at room temperature, under mixing in an end-to-end shaker. After this the activated MSNP-NH_2_ were collected and washed in PBS 1x thrice by centrifugation (1350g, 5 min), finally being re-suspended in a solution of urease (3 mg/mL) and heterobifunctional PEG (1 μg PEG /mg of MSNP-NH_2_) in PBS 1x. This mixture was left reacting in an end-to-end shaker, for 24 hours at room temperature. The resulting nanomotors were then collected and washed three times in PBS 1x by centrifugation (1350g, 5 min). Then, these nanomotors were re-suspended in a dispersion of AuNPs and left rotating in an end-to-end shaker for 10 minutes, followed by thorough washing by centrifugation (5 times, 1350g, 5 min).

### Dynamic Light Scattering and Electrophoretic Mobility Characterization of Nanomotors

A Wyatt Möbius DLS was used to characterized the hydrodynamic size distribution and surface charge of the MSNPs, MSNP-NH_2_, GA-activated MSNP-NH_2_, nanomotors and AuNPs-decorated nanomotors. The equipment comprises a 532 nm wavelength and a detector angle of 163.5° and it is able to analyze for light scattering and electrophoretic mobility simultaneously. We analyzed each of the particle types at a concentration of 20 μg/mL, with an acquisition time of 5 s and 3 runs per experiment. A total of 9 measurements per type of particle was performed to obtain statistically relevant data.

### Transmission Electron Microscopy (TEM) Imaging of the Nanomotors

The AuNPs-decorated nanomotors were diluted to a final concentration of 20 μg/mL in water, and the TEM images were captured.

### Synthesis of 6-[18F]Fluoronicotinic Acid 2,3,5,6-Tetrafluorophenyl Ester ([18F]F-PyTFP)

[^18^F]F-PyTFP was synthesized following a previously described procedure with modifications.(*66*) In brief, aqueous [^18^F]fluoride was first trapped in an ion-exchange resin (Sep-Pak^®^ Accell Plus QMA Light) and subsequently eluted to the reactor vessel with a solution of Kryptofix K_2.2.2_/K_2_CO_3_ in a mixture of water and acetonitrile. After azeotropic drying of the solvent, a solution of F-PyTFP (10 mg) in a mixture of tert-butanol and acetonitrile (4/1) was added and heated at 40°C for 15 min. The reaction mixture was then diluted with 1 mL of acetonitrile and 1 mL of water, and purified by HPLC using a Nucleosil 100-7 C18 column (Machery-Nagel, Düren, Germany) as stationary phase and 0.1%TFA/acetonitrile (25/75) as the mobile phase at a flow rate of 3 mL/min. The desired fraction (retention time = 22-23 minutes; [^18^F]F-PyTFP) was collected, diluted with water (25 mL), and flushed through a C18 cartridge (Sep-Pak^®^ Light, Waters) to selectively retain [^18^F]F-PyTFP. The desired labeled specie was finally eluted with acetonitrile (1 mL). Radiochemical purity was determined by radio-HPLC using a Mediterranean C18 column (4.6 × 150 mm, 5 μm) as stationary phase and 0.1% TFA/acetonitrile (0-1 minutes 25% acetonitrile; 9-12 minutes 90% acetonitrile; 13-15 minutes 25% acetonitrile) as the mobile phase at a flow rate of 1.5 mL/min (retention time =23 minutes).

### Radiolabelling of AuNPs-decorated nanomotors with [18F]F-PyTFP

The radiofluorination of urease-gold nanomotors with ^18^F was carried out by the reaction between the free amine groups of urease (e.g. present in lysine residues) contained at the surface of the nanomotors and [^18^F]F-PyTFP. Briefly, 200 μL of AuNPs-decorated nanomotors solution (1 mg/mL) were centrifuged during 10 minutes at 14800 rpm, re-suspended in PBS (10 μL, 10 mM pH 8) and mixed with 4 μL of [^18^F]F-PyTFP in ACN (approximately 74 MBq). The reaction mixture was incubated at room temperature for 35 min. After incubation, the crude was diluted with water (100 μL) and purified by centrifugation (5 minutes, 14800 rpm). The resulting precipitate was washed three times with water. The amount of radioactivity in the supernatant and the precipitate were determined in a dose calibrator (CPCRC-25R, Capintec Inc., NJ, USA) and analyzed with radio-thin layer chromatography (radio-TLC) using iTLC-SG chromatography paper (Agilent Technologies, CA, USA) and dichloromethane and methanol (2:1) as the stationary and mobile phases, respectively. TLC plates were analyzed using TLC-reader (MiniGITA, Raytest).

### Radiolabelling of BSA-MSNP with [18F]F-PyTFP

The radiofluorination of BSA-gold nanomotors with ^18^F was carried out following the same procedure described for AuNPs-decorated nanomotors above.

### Radiolabelling of AuNPs-decorated nanomotors with Iodine-124

The radioiodination of urease-gold nanomotors was performed by incubation with [^124^I]NaI. In brief, 200 μL of urease-gold nanomotors solution (1 mg/mL) diluted in 100 μL of water and 8 μL of [^124^I]NaI (approximately 1L MBq) were incubated at room temperature for 30 min. After incubation, the reaction mixture was purified by centrifugation (5 minutes, 14800 rpm). The resulting precipitate was washed three times with water (100 μL). The amount of radioactivity in the supernatant and the precipitate were determined in a dose calibrator (CPCRC-25R, Capintec Inc., NJ, USA) and analyzed with radio-thin layer chromatography (radio-TLC) using iTLC-SG chromatography paper (Agilent Technologies, CA, USA) and dichloromethane and methanol (2:1) as the stationary and mobile phases, respectively. TLC plates were analyzed using TLC-reader (MiniGITA, Raytest).

### Optical Video Recording

The optical videos of the swarms of nanomotors were acquired using a Leica DMi8 microscope, coupled with a Hamamatsu high-speed CCCD camera and a 2.5X objective. For this, the AuNPs-decorated nanomotors were centrifuged and re-suspended in 200 μL of PBS 1x. Then, a petri dish was filled with 3 mL of either PBS or a 300 mM solution of urea in PBS and placed in the microscope. A drop of 5 μL of the nanomotors was then added to the liquid-filled petri dish and 2 minute videos were acquired at a frame rate of 25 fps.

### Histogram Analysis

The acquired videos were then analyzed for pixel intensity distribution. For this, a region of interest (ROI) was selected, satisfying conditions such that it encloses both a part of the nanomotors population, but also a portion of the background, and a dimension of 300×300 pixels. Following this, the pixel intensity distribution within the ROI was analyzed at 12 s intervals, using ImageJ software.

### Density Map Analysis

The density maps of the optical videos were obtained using a custom made python code. Prior to this, the videos were treated to remove the background using ImageJ software, and the greyscale images were inverted. Then, the sum of pixel intensity of blocks of 1000 frames was calculated and the resulting images were loaded in the python code, applying the look-up table *turbo*.(*67*)

### Particle Image Velocimetry (PIV) Analysis

The particle image velocimetry analysis of the optical videos was performed using a custom made python code based on the OpenPIV library. Prior to loading the images in the code, the frames of the videos were treated to remove the background using ImageJ software. Following this, the resulting images were loaded in the python code, with a window size of 24 pixels, window overlap of 3 pixels, search size of 25 pixels and a frame rate of 0.4 s. Spurious vectors were removed by applying local, global, and signal-to-noise-ratio filters.

### Phantom Fabrication

To analyze the effect of complex paths on the motility of passive nanoparticles and active nanomotor swarms, phantoms with different shapes were fabricated. To obtain the desired channels’ geometry, a 3D design was prepared by using autoCAD software and post treated with PreForm Software to be later 3D printed by stereolitography. The rigid mold containing the inverse design of the desired channels was printed, followed by the required post-processing steps: (i) removal of the non-polymerized resin by two sequential washing steps in an isopropanol bath, (ii) hardening of the photopolymerized resin by 15 minutes of UV exposure, and (iii) removal of support structures. To fabricate the flexible and transparent PDMS-based channels, the catalyzer and the monomer were firstly mixed at a ratio of 1:10 and the solution was degassed for 15 minutes to avoid the bubble presence on the final chip. The solution was poured onto the rigid mold, followed by the curing process at 65°C overnight. The polymerized PDMS was then removed from the rigid mold, containing the desired channels. At this stage, the tubbing was implemented in the system, removing any debris that would fall into the channels. In order to close the system and obtain the phantom, a 2 mm thick layer of flat PDMS was bound to the open side of the channels using a 2 minute plasma treatment.

### PET-CT image acquisition

In vitro imaging studies were conducted using PET-CT as molecular imaging techniques. All the phantoms were filled either with 300 mM urea solution in water or ultrapure water (one channel with each medium) and positioned in the center of the field of view of the MOLECUBES β-CUBE (PET) scanner. The field of view was selected to cover the whole length of the phantom. For each type of nanomotor, two samples (10 μL, approximately 1 MBq each) were seeded simultaneously in one of the edges of the phantom, one in the channel filled with urea solution and the other one in the channel filled with ultrapure water. Immediately after, a dynamic PET scan was acquired for 25 minutes, followed by a CT acquisition.

### Phantom PET-CT imaging analysis

PET images were reconstructed and analyzed using PMOD image processing tool. With that aim, the whole channel was divided in sectors with the same length over the coronal view, and the concentration of activity in each section was determined as a function on time. The values of activity concentration were finally normalized to the whole amount of radioactivity in the channel.

### Intravenous administration

Anesthesia was induced by inhalation of 3% isoflurane in pure O_2_ and maintained by 1.5-2% isofluorane in 100% O_2_. With the animal under anesthesia, the F-nanomotors or I-nanomotors were injected via one of the lateral tail veins using PBS pH=7.4 as the vehicle (N=2 for each type of radiolabelling; see Table 1 for details). Dynamic, whole body 60-min PET imaging sessions were immediately started after administration of the labeled compounds using a MOLECUBES β-CUBE scanner. After the PET scan, whole body high resolution computerize tomography (CT) acquisitions were performed on the MOLECUBES X-CUBE scanner to provide anatomical information of each animal as well as the attenuation map for the later reconstruction of the PET images.

**Table 1.** Summary of the *in vivo* studies performed with the different types nanomotors and administration routes.

| Group | Number of animals | Particle | Administration route | Vehicle | Activity (MBq) | Mass of Particles (μg) |
| --- | --- | --- | --- | --- | --- | --- |
| 1 | 2 | F-nanomotors | Intravenous | PBS | 2.2±0.10 | 73 |
| 2 | 2 | I-nanomotors | Intravenous | PBS | 1.6±0.10 | 73 |
| 3 | 2 | F-nanomotors | intravesical | water | 1.6±0.10 | 33 |
| 4 | 2 | F- nanomotors | intravesical | urea | 1.9±0.15 | 33 |
| 5 | 2 | I-nanomotors | intravesical | water | 0.2±0.03 | 33 |
| 6 | 2 | I-nanomotors | intravesical | urea | 0.5±0.04 | 33 |
| 7 | 2 | MSNP-F-BSA | intravesical | water | 0.5±0.06 | 33 |
| 8 | 2 | MSNP-F-BSA | intravesical | urea | 0.4±0.05 | 33 |

### Intravesical administration

Anesthesia was induced by inhalation of 3% isoflurane in pure O_2_ and maintained by 1.5-2% isofluorane in 100% O_2_. With the animal under anesthesia, the animals were positioned in supine position and the bladder was emptied by massaging the abdominal region. Immediately after, the F-nanomotors and MSNP-F-BSA were introduced in the bladder (through a catheter) by intravesical administration using 300 mM of urea or water as the vehicle (N=2 for each type of radiolabeling and vehicle; see Table 1 for details). Administration was followed by 45-minutes PET imaging sessions and whole body high resolution CT acquisitions as above. PET images were reconstructed using 3D OSEM reconstruction algorithm and applying random, scatter and attenuation corrections. PET-CT images of the same mouse were co-registered and analyzed using PMOD image processing tool.

### Biodistribution analysis

Volumes of interest (VOIs) were placed on selected organs (namely: brain, thyroid, lungs, liver, stomach, kidneys, spleen, and bladder), as well as the heart in order to get an estimation of the concentration of radioactivity in blood. Time–activity curves (decay corrected) were obtained as cps/cm^3^ in each organ. Curves were transformed into real activity (Bq/cm^3^) curves by using a calibration factor, obtained from previous scans performed on a phantom (micro-deluxe, Data spectrum Corp.) under the same experimental conditions (isotope, reconstruction algorithm and energetic window).

### Bladder distribution analysis

PET images were reconstructed using 3D OSEM reconstruction algorithm and applying random, scatter and attenuation corrections. PET-CT images of the same mouse were co-registered and analyzed using PMOD image processing tool. Two VOIs were placed on the upper and lower regions of the bladder (namely: VOI 1 and VOI 2) to obtain the concentration of radioactivity in both VOIs over time. The values were normalized to the maximum values for each frame.

## Acknowledgments

The research leading to these results has received funding from the Spanish MINECO (BOTSinFluids project), the Foundation BBVA (MEDIROBOTS project), the CERCA program by the Generalitat de Catalunya and the CaixaImpulse program by La Caixa Foundation (TERANOBOTS project). A.C.H. thanks MINECO for the Severo Ochoa PhD fellowship. M.G. thanks MINECO for the Juan de la Cierva fellowship (IJCI2016-30451), the Beatriu de Pinós Programme (2018-BP-00305) and the Ministry of Business and Knowledge of the Government of Catalonia. D.V. acknowledges financial support provided by the European Commission under Horizon 2020s Marie Skłodowska-Curie Actions COFUND scheme [Grant Agreement No. 712754] and by the Severo Ochoa programme of the Spanish Ministry of Economy and Competitiveness [Grant SEV-2014-0425 (2015–2019)]. T.P. thanks the European Union’s Horizon 2020 research and innovation program, under the Marie Skłodowska-Curie Individual Fellowship (H2020-MSCA-IF2018, DNA-bots). J.L. thanks The Spanish Ministry of Economy and Competitiveness (Grant CTQ2017-87637-R) for financial support. Part of the work was conducted under the Maria de Maeztu Units of Excellence Programme – Grant No. MDM-2017-0720.

## Author contributions

A.C.H synthesized the nanomotors, performed their characterization, optical tracking, corresponding image processing and data analysis; C.S. performed radiolabeling, PET-CT experiments and data analysis. M.G. fabricated the phantoms and helped with graphic design and video editing. S. G.-G. and E.J. helped set-up the intravesical administration model. D.V. synthesized the AuNPs and performed their characterization. L.R developed the ^18^F-radiolabeling protocol. P.R.-C provided graphic design of the radiochemical facility and contributed in image analysis after intravesical instillation. U.C. performed reconstruction, analysis and quantification of PET images. V.G.-V contributed to the *in vivo* experimental design. T.P., J.L and S.S. conceived the idea, designed and supervised the work. The manuscript was written through contributions of all authors. All authors have given approval to the final version of the manuscript.

## Data and materials availability

Supplementary information is available for this paper online.

## Supporting Information

### Supplementary Figures

**Figure S1.**
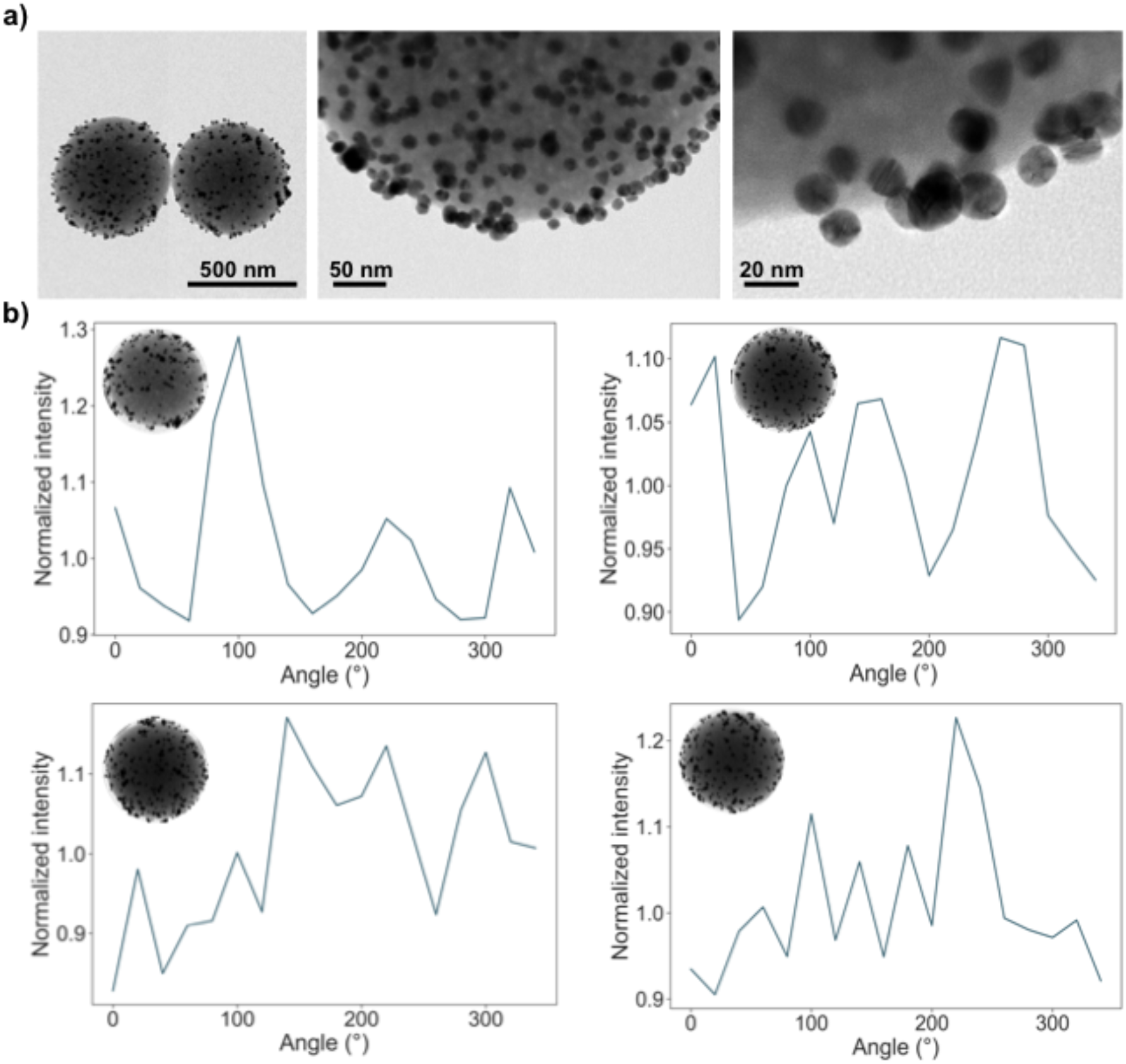
Characterization of the AuNP-decorated nanomotors. a) Transmission electron microscopy images. b) Analysis of the radial pixel intensity to demonstrate the asymmetry of AuNPs distribution on the surface of the nanomotors.

**Figure S2.**
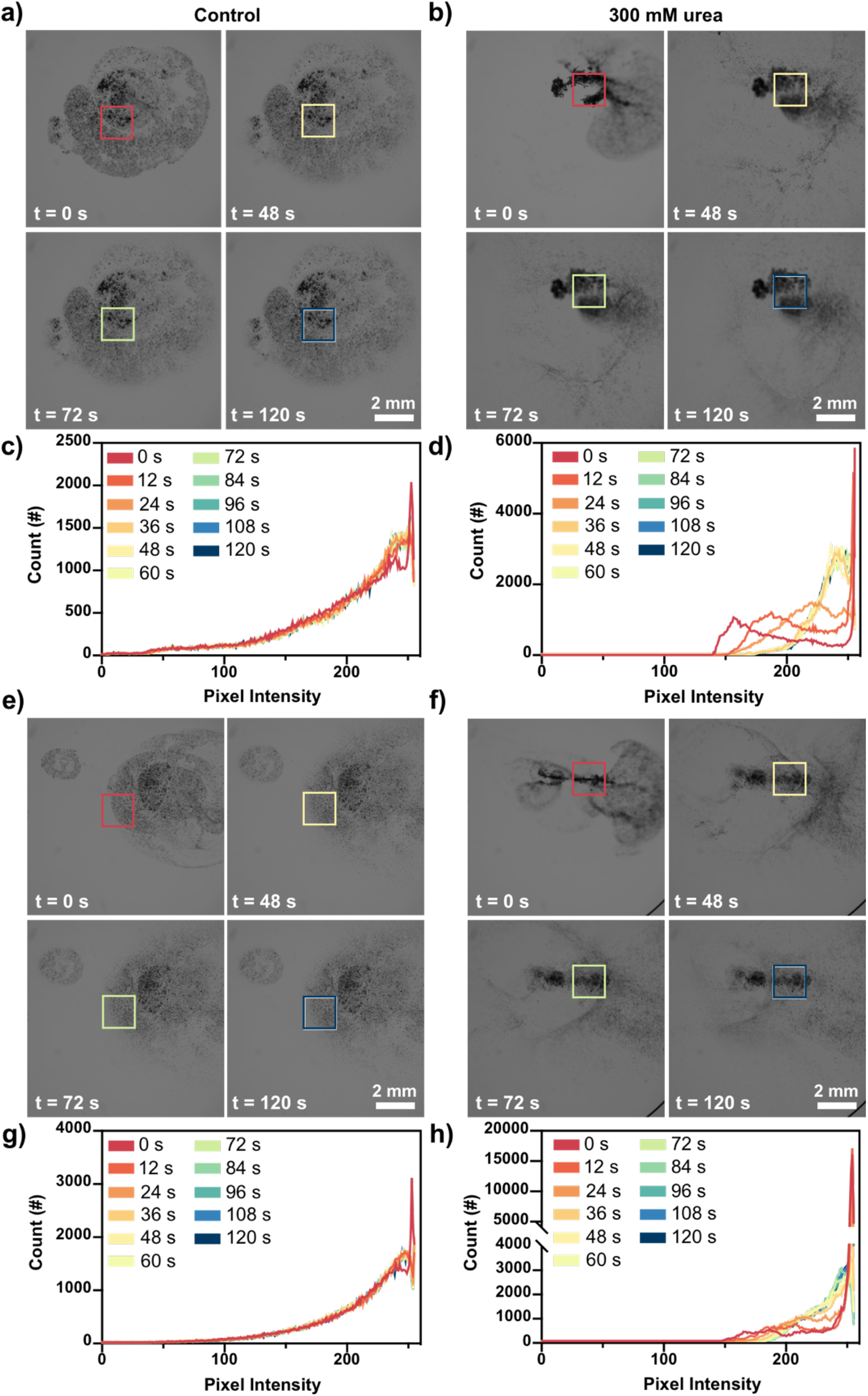
In vitro analysis of the motion of swarms of urease-powered AuNP decorated nanomotors both by optical microscopy. Snapshots of a population of urease-powered nanomotors in a bath of PBS (a and e), imaged by optical microscopy. Snapshots of a population of urease-powered nanomotors in a bath of 300 mM urea in PBS (b and f), imaged by optical microscopy. Pixel intensity distribution histograms on a region of interest of the population of nanomotors in a bath of PBS (c and g). Pixel intensity distribution histograms on a region of interest of the population of nanomotors in a bath of 300 mM urea in PBS (d and h).

**Figure S3.**
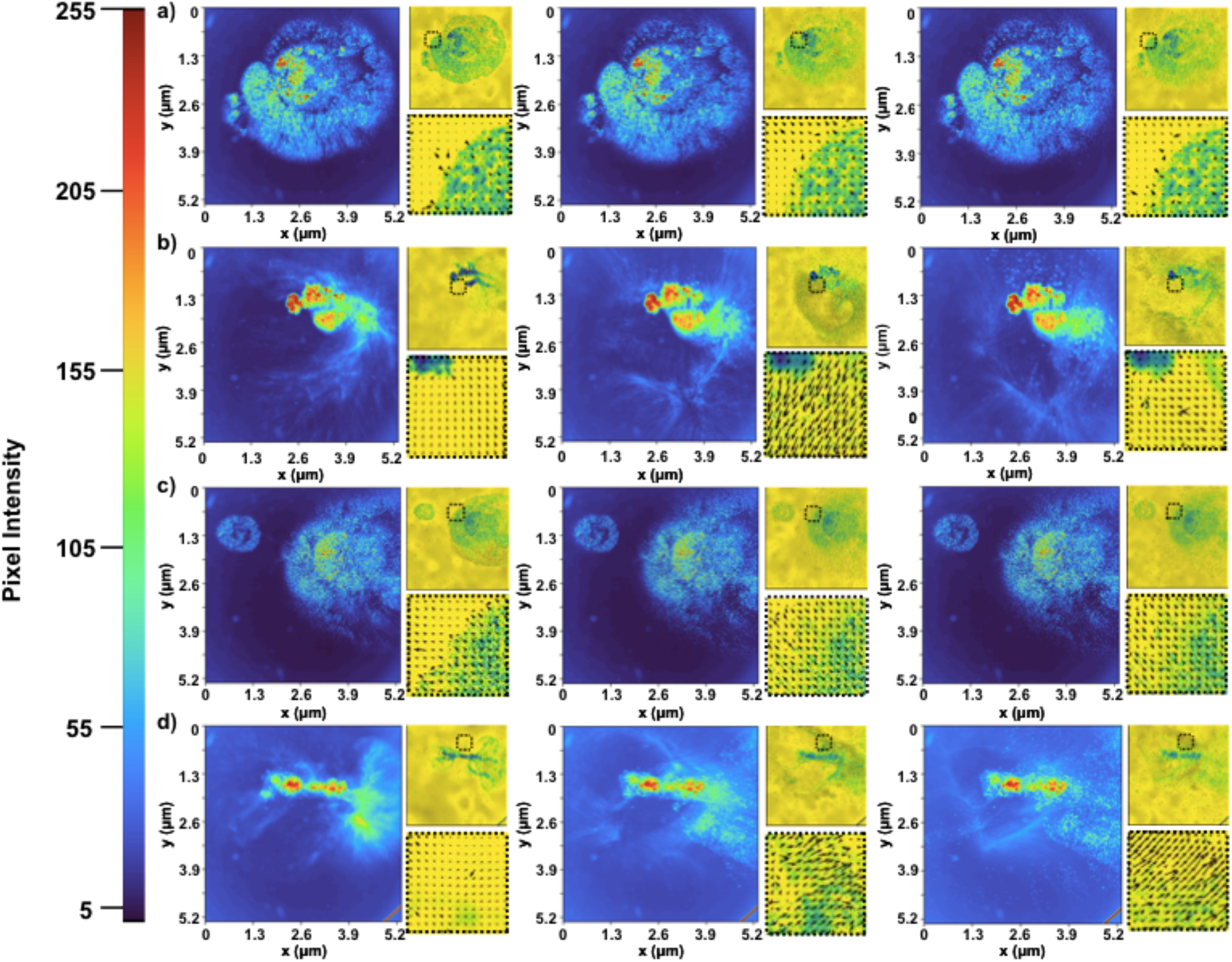
Motion analysis of the nanomotor swarms by optical microscopy. Density maps obtained by calculating the average pixel intensity over periods of 180 seconds of the videos and particle image velocimetry (PIV) of the videos of nanomotors in a bath of PBS (a and c) and nanomotors in a bath of 300 mM urea in PBS (b and d).

**Figure S4.**
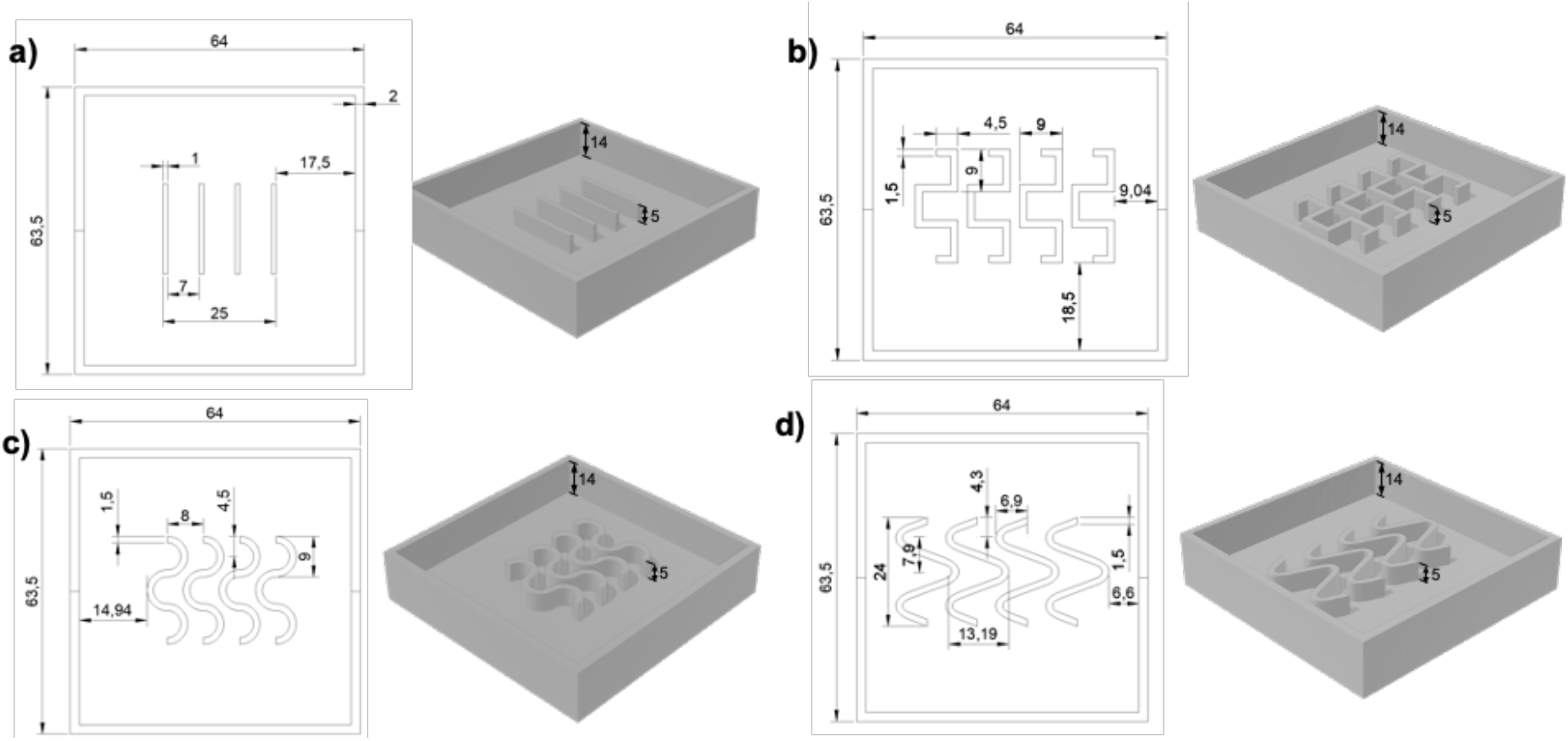
Fabrication of the molds for the complex paths phantoms. Design and dimensions of a) mold of phantom i, b) mold of phantom ii, c) mold of phantom iii and d) mold of phantom iv.

**Figure S5.**
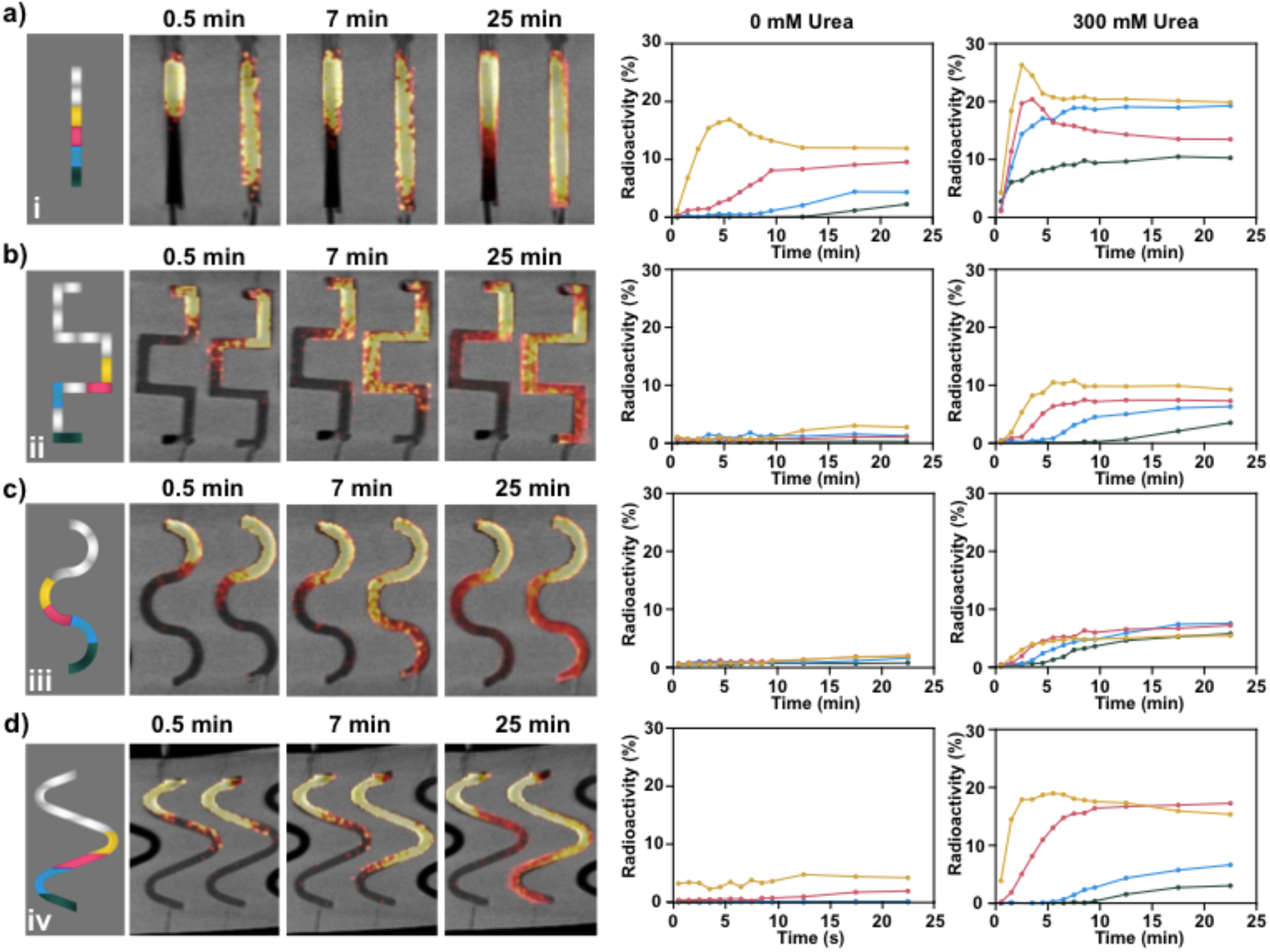
PET-CT coronal images obtained a 0.5 minutes, 7 minutes and 25 minutes for each phantom, which was filled either with Milli-Q water or 300 mM urea solution. The channels were then divided in sectors with the same length over the coronal view and the concentration of activity in each channel was measured as a function on time. The values were normalized to the total amount of radioactivity in the channel. The snapshots of coronal view and analysis of sectors are depicted in a) for the straight channels phantom (I), b) for the squared phantom (ii), c) for the curved channels phantom (iii) and d) for the more curved channels phantom (iv). e) Percentage of radioactivity in the sector A of each phantom at the end of the experiment. f) Sum of the percentage of radioactivity in the second half of each phantom at the end of the experiment and corresponding fold increase in radioactivity due to the presence of urea.

**Figure S6.**
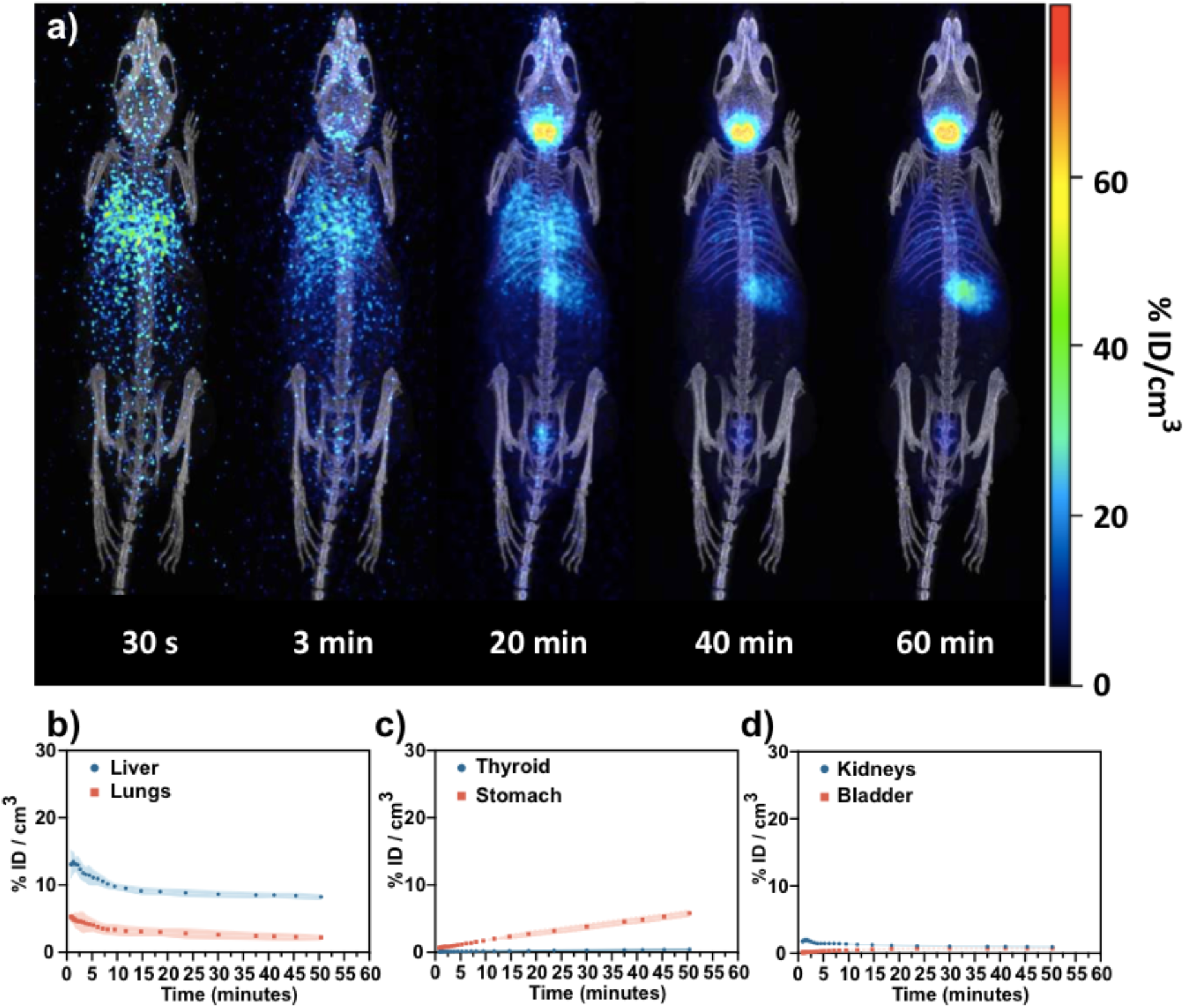
Analysis of the biodistribution of I-nanomotors injected intravenously. a) PET-CT images (maximum intensity projections, coronal views) obtained at different time points after intravenous administration of I-nanomotors. Accumulation of I-nanomotors in b) liver and lungs and c) thyroid and stomach, and d) kidneys and bladder, as determined by PET imaging. Results are expressed as % of injected dose per organ. Error bars correspond to the SD (N = 2)

### List of Supplementary Videos

**Video S1.** In vitro imaging of enzyme-powered nanomotor swarms by optical microscopy.

**Video S2.** In vitro imaging of enzyme-powered nanomotor swarms by PET.

**Video S3.** Effect of complex paths on the *in vitro* motion of ^18^F radiolabeled urease-powered nanomotors studied by PET-CT imaging.

**Video S4.** Study of the biodistribution of enzyme-powered nanomotors by PET-CT.

**Video S5.** PET-CT analysis of the distribution of 18F-nanomotors and MSNP-18F-BSA in the bladder administered via intravesical instillation.

## References

1. S. Wilhelm, A. J. Tavares, Q. Dai, S. Ohta, J. Audet, H. F. Dvorak, W. C. W. Chan, Analysis of nanoparticle delivery to tumours. Nat. Rev. Mater. 1, 16014 (2016).

2. Z. Wu, X. Lin, Y. Wu, T. Si, J. Sun, Q. He, Near-Infrared Light-Triggered “On/Off” Motion of Polymer Multilayer Rockets. ACS Nano. 8, 6097–6105 (2014).

3. S. M. Douglas, I. Bachelet, G. M. Church, A Logic-Gated Nanorobot for Targeted Transport of Molecular Payloads. Science. 335, 831–834 (2012).

4. A. C. Hortelao, R. Carrascosa, N. Murillo-Cremaes, T. Patino, S. Sánchez, Targeting 3D Bladder Cancer Spheroids with Urease-Powered Nanomotors. ACS Nano. 13, 429–439 (2019).

5. F. Hallouard, N. Anton, P. Choquet, A. Constantinesco, T. Vandamme, Iodinated blood pool contrast media for preclinical X-ray imaging applications--a review. Biomaterials. 31, 6249–6268 (2010).

6. A. C. Hortelão, T. Patiño, A. Perez-Jiménez, À. Blanco, S. Sánchez, Enzyme-Powered Nanobots Enhance Anticancer Drug Delivery. Adv. Funct. Mater. 28, 1705086 (2018).

7. F. Peng, Y. Tu, D. A. Wilson, Micro/nanomotors towards in vivo application: cell, tissue and biofluid. Chem. Soc. Rev. 46, 5289–5310 (2017).

8. A. Llopis-Lorente, A. Garciá-Fernández, N. Murillo-Cremaes, A. C. Hortelaõ, T. Patinõ, R. Villalonga, F. Sancenón, R. Martínez-Máñez, S. Sánchez, Enzyme-powered gated mesoporous silica nanomotors for on-command intracellular payload delivery. ACS Nano. 13, 12171–12183 (2019).

9. S. Tang, F. Zhang, H. Gong, F. Wei, J. Zhuang, E. Karshalev, B. Esteban-Fernández de Ávila, C. Huang, Z. Zhou, Z. Li, L. Yin, H. Dong, R. H. Fang, X. Zhang, L. Zhang, J. Wang, Enzyme-powered Janus platelet cell robots for active and targeted drug delivery. Sci. Robot. 5, eaba6137 (2020).

10. W. Gao, R. Dong, S. Thamphiwatana, J. Li, W. Gao, L. Zhang, J. Wang, Artificial Micromotors in the Mouse’s Stomach: A Step toward in Vivo Use of Synthetic Motors. ACS Nano. 9, 117–123 (2015).

11. D. Schamel, A. G. Mark, J. G. Gibbs, C. Miksch, K. I. Morozov, A. M. Leshansky, P. Fischer, Nanopropellers and Their Actuation in Complex Viscoelastic Media. ACS Nano. 8, 8794–8801 (2014).

12. H. Choi, S. H. Cho, S. K. Hahn, Urease-Powered Polydopamine Nanomotors for Intravesical Therapy of Bladder Diseases. ACS Nano (2020), doi:10.1021/acsnano.9b09726.

13. D. Walker, B. T. Kasdorf, H.-H. Jeong, O. Lieleg, P. Fischer, Enzymatically active biomimetic micropropellers for the penetration of mucin gels. Sci. Adv. 1, e1500501–e1500501 (2015).

14. J. Wang, B. J. Toebes, A. S. Plachokova, Q. Liu, D. Deng, J. A. Jansen, F. Yang, D. A. Wilson, Self-Propelled PLGA Micromotor with Chemotactic Response to Inflammation. Adv. Healthc. Mater. 9, 1901710 (2020).

15. F. Peng, Y. Tu, A. Adhikari, J. C. J. Hintzen, D. W. P. M. Lowik, D. A. Wilson, Peptide functionalized nanomotor as efficient cell penetrating tool. Chem. Commun. 53, 1088–1091 (2016).

16. J. Sun, M. Mathesh, W. Li, D. A. Wilson, Enzyme-Powered Nanomotors with Controlled Size for Biomedical Applications. ACS Nano. 13, 10191–10200 (2019).

17. M. A. Ramos-Docampo, M. Fernández-Medina, E. Taipaleenmäki, O. Hovorka, V. Salgueiriño, B. Städler, Microswimmers with Heat Delivery Capacity for 3D Cell Spheroid Penetration. ACS Nano. 13, 12192–12205 (2019).

18. G. von Maltzahn, J.-H. Park, K. Y. Lin, N. Singh, C. Schwöppe, R. Mesters, W. E. Berdel, E. Ruoslahti, M. J. Sailor, S. N. Bhatia, Nanoparticles that communicate in vivo to amplify tumour targeting. Nat. Mater. 10, 545–552 (2011).

19. D. Needleman, Z. Dogic, Active matter at the interface between materials science and cell biology. Nat. Rev. Mater. 2, 17048 (2017).

20. M. Whiteley, S. P. Diggle, E. P. Greenberg, Progress in and promise of bacterial quorum sensing research. Nature. 551, 313–320 (2017).

21. K. Kawabata, M. Nagayama, H. Haga, T. Sambongi, Mechanical effects on collective phenomena of biological systems: cell locomotion. Curr. Appl. Phys. 1, 66–71 (2001).

22. Z.-M. Qian, S. H. Wang, X. E. Cheng, Y. Q. Chen, An effective and robust method for tracking multiple fish in video image based on fish head detection. BMC Bioinformatics. 17, 251 (2016).

23. Z. Deng, F. Mou, S. Tang, L. Xu, M. Luo, J. Guan, Swarming and collective migration of micromotors under near infrared light. Appl. Mater. Today. 13, 45–53 (2018).

24. X. Yan, Q. Zhou, M. Vincent, Y. Deng, J. Yu, J. Xu, T. Xu, T. Tang, L. Bian, Y.-X. J. Wang, K. Kostarelos, L. Zhang, Multifunctional biohybrid magnetite microrobots for imaging-guided therapy. Sci. Robot. 2, eaaq1155 (2017).

25. A. Somasundar, S. Ghosh, F. Mohajerani, L. N. Massenburg, T. Yang, P. S. Cremer, D. Velegol, A. Sen, Positive and negative chemotaxis of enzyme-coated liposome motors. Nat. Nanotechnol. 14(2019), doi:10.1038/s41565-019-0578-8.

26. Z. Wu, L. Li, Y. Yang, P. Hu, Y. Li, S.-Y. Yang, L. V Wang, W. Gao, A microrobotic system guided by photoacoustic computed tomography for targeted navigation in intestines in vivo. Sci. Robot. 4, eaax0613 (2019).

27. A. Aubret, M. Youssef, S. Sacanna, J. Palacci, Targeted assembly and synchronization of self-spinning microgears. Nat. Phys. 14, 1114–1118 (2018).

28. C. Zhou, N. J. Suematsu, Y. Peng, Q. Wang, X. Chen, Y. Gao, W. Wang, Coordinating an Ensemble of Chemical Micromotors via Spontaneous Synchronization. ACS Nano (2020), doi:10.1021/acsnano.9b08421.

29. X. Chen, C. Zhou, Y. Peng, Q. Wang, W. Wang, Temporal Light Modulation of Photochemically Active, Oscillating Micromotors: Dark Pulses, Mode Switching, and Controlled Clustering. ACS Appl. Mater. Interfaces. 12, 11843–11851 (2020).

30. X. Cao, E. Panizon, A. Vanossi, N. Manini, C. Bechinger, Orientational and directional locking of colloidal clusters driven across periodic surfaces. Nat. Phys. 15, 776–780 (2019).

31. D. Zhou, Y. Gao, J. Yang, Y. C. Li, G. Shao, G. Zhang, T. Li, L. Li, Light-Ultrasound Driven Collective “Firework” Behavior of Nanomotors. Adv. Sci. 5, 1800122 (2018).

32. F. Mou, X. Li, Q. Xie, J. Zhang, K. Xiong, L. Xu, J. Guan, Active Micromotor Systems Built from Passive Particles with Biomimetic Predator–Prey Interactions. ACS Nano. 14, 406–414 (2020).

33. T. Xu, F. Soto, W. Gao, R. Dong, V. Garcia-Gradilla, E. Magaña, X. Zhang, J. Wang, Reversible swarming and separation of self-propelled chemically powered nanomotors under acoustic fields. J. Am. Chem. Soc. 137, 2163–2166 (2015).

34. M. Ibele, T. E. Mallouk, A. Sen, Schooling Behavior of Light-Powered Autonomous Micromotors in Water. Angew. Chemie Int. Ed. 48, 3308–3312 (2009).

35. J. Yan, M. Han, J. Zhang, C. Xu, E. Luijten, S. Granick, Reconfiguring active particles by electrostatic imbalance. Nat. Mater. 15, 1095–1099 (2016).

36. I. Buttinoni, J. Bialké, F. Kümmel, H. Löwen, C. Bechinger, T. Speck, Dynamical Clustering and Phase Separation in Suspensions of Self-Propelled Colloidal Particles. Phys. Rev. Lett. 110, 238301 (2013).

37. T. Bäuerle, A. Fischer, T. Speck, C. Bechinger, Self-organization of active particles by quorum sensing rules. Nat. Commun. 9, 3232 (2018).

38. J. Palacci, S. Sacanna, A. P. Steinberg, D. J. Pine, P. M. Chaikin, Living Crystals of Light-Activated Colloidal Surfers. Science. 339, 936–940 (2013).

39. J. Yu, D. Jin, K.-F. Chan, Q. Wang, K. Yuan, L. Zhang, Active generation and magnetic actuation of microrobotic swarms in bio-fluids. Nat. Commun. 10, 5631 (2019).

40. Z. Wu, J. Troll, H.-H. Jeong, Q. Wei, M. Stang, F. Ziemssen, Z. Wang, M. Dong, S. Schnichels, T. Qiu, P. Fischer, A swarm of slippery micropropellers penetrates the vitreous body of the eye. Sci. Adv. 4, eaat4388 (2018).

41. A. Servant, F. Qiu, M. Mazza, K. Kostarelos, B. J. Nelson, Controlled In Vivo Swimming of a Swarm of Bacteria-Like Microrobotic Flagella. Adv. Mater. 27, 2981–2988 (2015).

42. M. Bauer, C. C. Wagner, O. Langer, Microdosing Studies in Humans. Drugs R D. 9, 73–81 (2008).

43. M. Bergstrom, The Use of Microdosing in the Development of Small Organic and Protein Therapeutics. J. Nucl. Med. . 58, 1188–1195 (2017).

44. D. Vilela, U. Cossío, J. Parmar, A. M. Martínez-Villacorta, V. Gómez-Vallejo, J. Llop, S. Sánchez, Medical Imaging for the Tracking of Micromotors. ACS Nano. 12, 1220–1227 (2018).

45. F. Tang, L. Li, D. Chen, Mesoporous Silica Nanoparticles: Synthesis, Biocompatibility and Drug Delivery. Adv. Mater. 24, 1504–1534 (2012).

46. Q. He, X. Cui, F. Cui, L. Guo, J. Shi, Size-controlled synthesis of monodispersed mesoporous silica nano-spheres under a neutral condition. Microporous Mesoporous Mater. 117, 609–616 (2009).

47. Y. Wang, Y. Sun, J. Wang, Y. Yang, Y. Li, Y. Yuan, C. Liu, Charge-Reversal APTES-Modified Mesoporous Silica Nanoparticles with High Drug Loading and Release Controllability. ACS Appl. Mater. Interfaces. 8, 17166–17175 (2016).

48. H. Xu, F. Yan, E. E. Monson, R. Kopelman, Room-temperature preparation and characterization of poly (ethylene glycol)-coated silica nanoparticles for biomedical applications. J. Biomed. Mater. Res. Part A. 66A, 870–879 (2003).

49. M. R. Ivanov, H. R. Bednar, A. J. Haes, Investigations of the Mechanism of Gold Nanoparticle Stability and Surface Functionalization in Capillary Electrophoresis. ACS Nano. 3, 386–394 (2009).

50. Z. Liu, Y. Yan, F. T. Chin, F. Wang, X. Chen, Dual Integrin and Gastrin-Releasing Peptide Receptor Targeted Tumor Imaging Using 18F-labeled PEGylated RGD-Bombesin Heterodimer 18F-FB-PEG3-Glu-RGD-BBN. J. Med. Chem. 52, 425–432 (2009).

51. K. R. Pulagam, K. B. Gona, V. Gómez-Vallejo, J. Meijer, C. Zilberfain, I. Estrela-Lopis, Z. Baz, U. Cossío, J. Llop, Gold nanoparticles as boron carriers for boron neutron capture therapy: Synthesis, radiolabelling and in vivo evaluation. Molecules. 24(2019), doi:10.3390/molecules24193609.

52. X. Ma, A. Jannasch, U.-R. Albrecht, K. Hahn, A. Miguel-López, E. Schäffer, S. Sánchez, Enzyme-Powered Hollow Mesoporous Janus Nanomotors. Nano Lett. 15, 7043–7050 (2015).

53. X. Arqué, A. Romero-Rivera, F. Feixas, T. Patiño, S. Osuna, S. Sánchez, Intrinsic enzymatic properties modulate the self-propulsion of micromotors. Nat. Commun. 10, 1–12 (2019).

54. T. Patiño, N. Feiner-Gracia, X. Arqué, A. Miguel-López, A. Jannasch, T. Stumpp, E. Schäffer, L. Albertazzi, S. Sánchez, Influence of Enzyme Quantity and Distribution on the Self-Propulsion of Non-Janus Urease-Powered Micromotors. J. Am. Chem. Soc. 140, 7896–7903 (2018).

55. M. De Corato, X. Arqué, T. Patiño, M. Arroyo, S. Sánchez, I. Pagonabarraga, Self-Propulsion of Active Colloids via Ion Release: Theory and Experiments. Phys. Rev. Lett. 124, 108001 (2020).

56. L. Liu, H. Mo, S. Wei, D. Raftery, Quantitative analysis of urea in human urine and serum by 1H nuclear magnetic resonance. Analyst. 137, 595–600 (2012).

57. L. Soler, V. Magdanz, V. M. Fomin, S. Sanchez, O. G. Schmidt, Self-Propelled Micromotors for Cleaning Polluted Water. ACS Nano. 7, 9611–9620 (2013).

58. J. Orozco, B. Jurado-Sánchez, G. Wagner, W. Gao, R. Vazquez-Duhalt, S. Sattayasamitsathit, M. Galarnyk, A. Cortés, D. Saintillan, J. Wang, Bubble-Propelled Micromotors for Enhanced Transport of Passive Tracers. Langmuir. 30, 5082–5087 (2014).

59. E. Morales-Narváez, M. Guix, M. Medina-Sánchez, C. C. Mayorga-Martinez, A. Merkoçi, Micromotor Enhanced Microarray Technology for Protein Detection. Small. 10, 2542–2548 (2014).

60. H. W. A. M. de Jong, L. Perk, G. W. M. Visser, R. Boellaard, G. A. M. S. van Dongen, A. A. Lammertsma, in IEEE Nuclear Science Symposium Conference Record, 2005 (2005), vol. 3, pp. 1624–1627.

61. N. Hoshyar, S. Gray, H. Han, G. Bao, The effect of nanoparticle size on in vivo pharmacokinetics and cellular interaction. Nanomedicine. 11, 673–692 (2016).

62. B. Du, X. Jiang, A. Das, Q. Zhou, M. Yu, R. Jin, J. Zheng, Glomerular barrier behaves as an atomically precise bandpass filter in a sub-nanometre regime. Nat. Nanotechnol. 12, 1096–1102 (2017).

63. S. Il Kim, S. H. Choo, in Bladder Cancer, J. H. Ku, Ed. (Elsevier, ed. 1st, 2018;), pp. 263–276.

64. K. Ai, Y. Liu, L. Lu, Hydrogen-Bonding Recognition-Induced Color Change of Gold Nanoparticles for Visual Detection of Melamine in Raw Milk and Infant Formula. J. Am. Chem. Soc. 131, 9496–9497 (2009).

65. D. Vilela, A. C. Hortelao, R. Balderas-Xicohténcatl, M. Hirscher, K. Hahn, X. Ma, S. Sánchez, Facile fabrication of mesoporous silica micro-jets with multi-functionalities. Nanoscale. 9, 13990–13997 (2017).

66. D. E. Olberg, J. M. Arukwe, D. Grace, O. K. Hjelstuen, M. Solbakken, G. M. Kindberg, Cuthbertson, One step radiosynthesis of 6-[18F]fluoronicotinic acid 2,3,5,6-tetrafluorophenyl ester ([18F]F-Py-TFP): A new prosthetic group for efficient labeling of biomolecules with fluorine-18. J. Med. Chem. (2010), doi:10.1021/jm9015813.

67. A. Mikhailov, Turbo, An Improved Rainbow Colormap for Visualization. Google AI Blog (2019).

